# Mitotic heritability of DNA methylation at intermediately methylated sites is imprecise

**DOI:** 10.1101/2023.01.27.525699

**Authors:** Amir D. Hay, Noah J. Kessler, Daniel Gebert, Nozomi Takahashi, Hugo Tavares, Felipe K. Teixeira, Anne C. Ferguson-Smith

## Abstract

DNA methylation is considered a stable epigenetic mark due to its presumed long-term inheritance through cell divisions. Here, we perform high-throughput bisulfite sequencing on clonally derived cell lines to quantitatively measure mitotic methylation inheritance at the nucleotide level. We find that although DNA methylation is generally faithfully maintained at hypo- and hypermethylated sites, this is not the case at intermediately methylated CpGs. Low fidelity intermediate methylation is interspersed throughout the genome and within genes with no or low transcriptional activity. Moreover, we determine that the probabilistic changes that occur at intermediately methylated sites are due to DNMT1 rather than DNMT3A/3B activity. The observed lack of clonal inheritance at intermediately methylated sites challenges the concept of DNA methylation as a consistently stable epigenetic mark.

---

The establishment and maintenance of global DNA methylation patterns are essential for development and function of vertebrate genomes (*1, 2*). However, accumulating evidence indicates that at a given CpG dinucleotide, the DNA methylation status may vary between cell divisions, suggesting that DNA methylation patterns are more dynamic than previously anticipated. One mechanism that can explain this is the dynamic binding of factors and chromatin states that modulate methylation deposition during development (*3*). In contrast, some early investigations found that intermediately methylated sites could spontaneously arise within subclonal cell populations derived from single cells (*4, 5*). Other groups observed that intermediate methylation is inconsistently inherited after cell divisions and therefore represents either errors in maintenance or spontaneous *de novo* methylation in a range of developmental and tumour cell populations (*6-8*). While models of how intermediate methylation may be inherited through cellular divisions have been proposed (*9, 10*), the extent of such dynamics has not been systematically examined at a genome-wide scale, nor have the principles dictating DNA methylation stability through clonal propagation been determined.

We devised an experiment to directly assess the fidelity of DNA methylation through cell divisions at the genome-scale, by subcloning populations of cells and performing target DNA capture followed by high-throughput bisulfite sequencing (tcBS-seq, see methods) on both the subclones and the parent population of cells (Fig. 1A). To do so, we established mouse embryonic fibroblasts (MEFs) from two sibling E13.5 mouse embryos and immortalised the cell lines. From these parental lines, we randomly sampled fourteen single cells to establish clonal populations of around 1-2 million cells on which we profiled DNA methylation using tcBS-seq. In total, the methylation levels of >1.2 million CpGs (or around 5% of CpGs in the mouse genome, with a median coverage of 32x per CpG per dataset; fig. S1) were determined across 16 samples and used for downstream analysis. In principle, in a single cell at a single CpG site, there are three possible stable (strand-symmetric) methylation states (Fig. 1B), which provides us with a tractable framework to determine the fidelity of DNA methylation inheritance through cell divisions. In a purely faithful scenario, the DNA methylation status of each CpG in the clonal lines should be exclusively either 0%, 50%, or 100%.

**Fig. 1.**
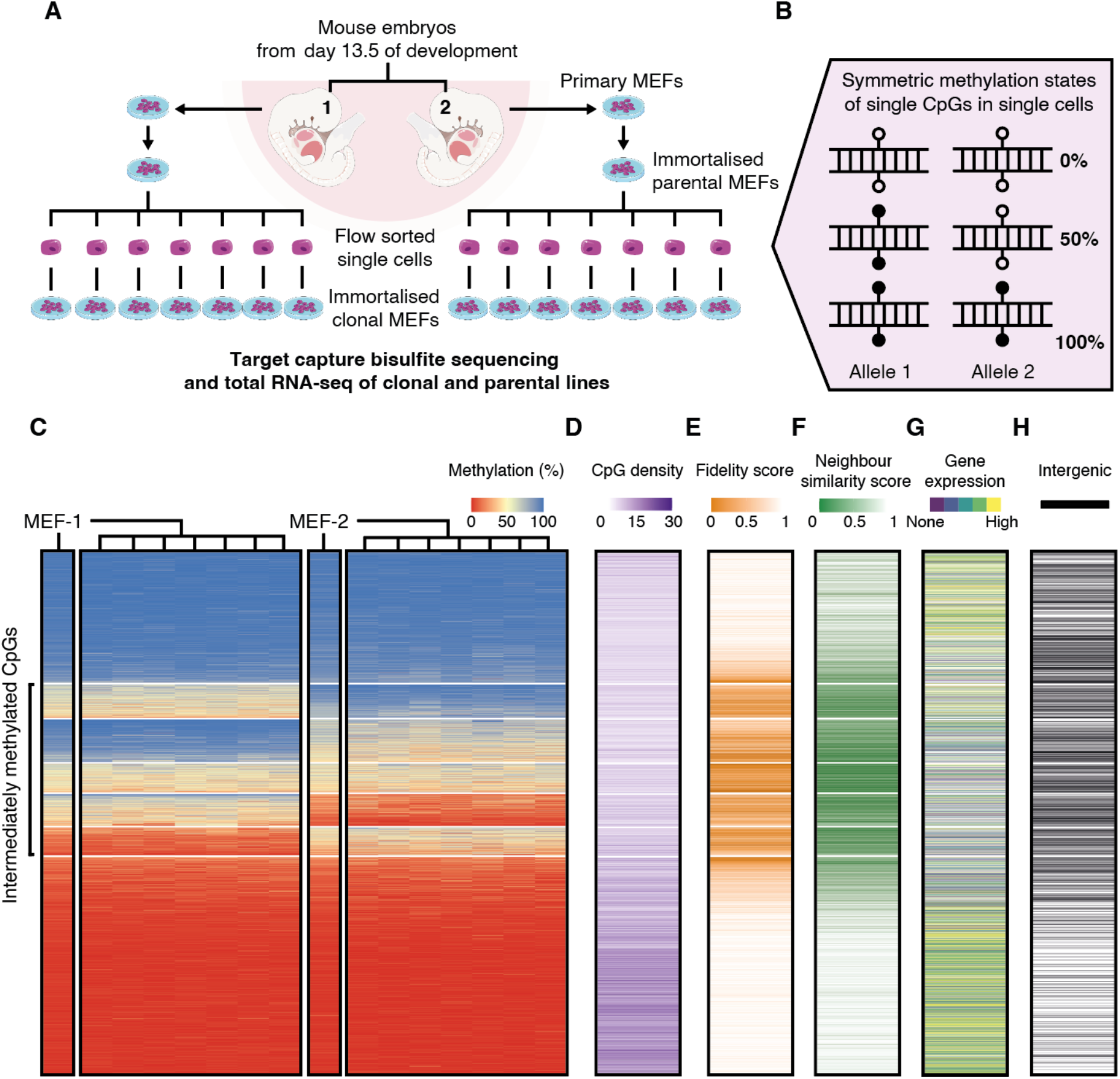
Intermediately methylated sites show lower epigenetic inheritance fidelity. (A) Mouse embryonic fibroblasts (MEFs) were isolated from E13.5 mouse embryos and immortalised. The resulting cell lines, MEF-1 and MEF-2, are referred to as the “parental” lines. Single cells were randomly selected by flow cytometry from these lines and grown into derivative “clonal” lines. Target capture bisulfite sequencing and total RNA sequencing were performed on both the parental and clonal lines. (B) Illustration showing that at the single-cell level, there are only three strand-symmetric methylation states that can exist at a single CpG dinucleotide: 0%, 50%, and 100%. (C) Heatmaps of 1,203,687 CpGs sorted by median methylation (%) within k-means clusters (k=7, separated by white lines). The two parental lines are denoted as MEF-1 and MEF-2, with the seven clonal lines shown to the right of the corresponding parental line. (D-H) Heatmaps of (D) CpG density calculated as the number of CpGs within 100bp of each focal CpG, (E) methylation fidelity score calculated as a proxy for the retention of symmetric methylation states from the single cell to a clonal line population, (F) neighbour similarity score as an approximation of the consistency of methylation between neighbouring CpGs in a clonal line, (G) and transcription quintiles derived from gene expression averages across all the cell lines. (H) Intergenic CpGs (blank lines) are characterised by a lack of overlap with an annotated protein coding transcript or promoter.

Globally, methylation data generated from parental and clonal lines revealed that the methylation state of most CpGs (66%) is consistent across both parental lines and all 14 clonal samples (Fig. 1C, fig. S1 and S2, and table S1 and S2). When CpGs are classified according to their methylation states using k-means clustering (fig. S3, A and B), we find that 40% of all tested CpGs are consistently hypomethylated (U) across all analysed lines and are enriched for CpG dense regions (Fig. 1D and fig. S3C). Conversely, 26% of CpGs are consistently hypermethylated (M). As expected, the frequency of these two states indicates that the majority are explained by faithful inheritance of DNA methylation through cell divisions. In agreement, CpGs that are consistently hypo/hypermethylated in MEF-1-derived lines, exhibit the same state in MEF-2-derived lines. On the other hand, 33% of CpGs are shown to be intermediately methylated across all clonal lines derived from at least one of the parental cell lines. Among these intermediately methylated CpGs, most (26% of all analysed CpGs) are consistently hypo/hypermethylated across one set of lines but intermediately methylated in the other set (UI or MI, respectively). In addition, a subset of CpGs (7%) was intermediately methylated across all the cell lines (I).

To determine the degree of faithful methylation inheritance at the CpG level, we calculated the fraction of clonal lines that exhibit 0%, 50% or 100% methylation. We refer to this metric as the fidelity score, based on the concept that faithful transmission of the methylation state would result in only 0%, 50%, or 100% methylation in the clonal line populations derived from single cells. As expected from the consistency of methylation states across the samples, we find that CpGs classified as M and U generally have a high fidelity score. On the other hand, CpGs that have the potential to be intermediately methylated have a significantly lower fidelity score (Fig 1E and fig S3D). Therefore, intermediate methylation states generally represent unfaithful inheritance of methylation between cell divisions.

To gain insight into the principles dictating DNA methylation stability, with the basis that intermediate levels reflect unfaithful inheritance between cell divisions, we compared the methylation groups with respect to their genomic sequence context. Because CpG density is a major predictor of methylation status (*11*), we first asked how CpG density correlates with the different methylation groups. As expected for consistently hypomethylated CpGs (*12*), we found that U CpGs reside in CpG dense regions in comparison to the other groups. On the other hand, M CpGs, as well as CpGs that have potential to be intermediately methylated (UI, I, MI), have significantly lower CpG density (Fig. 1D and fig. S3C). Given that methylation can be spatially regulated across multiple neighbouring CpGs (*13*), we calculated a methylation co-variation score between each one of the 1.2M CpGs and its closest neighbour (neighbour similarity score, see methods). We found that, compared to CpGs classified as U and M, CpGs with potential for intermediate methylation (UI, I, MI) are less likely to have a similar methylation level to the closest CpG (Fig. 1F and fig. S3E). Together, these results show that intermediately methylated CpGs, which are generally unfaithfully propagated, are enriched in regions of low CpG density, and attain methylation independently of neighbouring CpGs.

Since DNA methylation is associated with transcriptional repression (*14*), we performed total RNA-sequencing on the parental and clonal MEF lines to investigate whether there is a relationship between the fidelity of methylation and gene expression. First, we classified all protein-coding genes (21,835) into five expression level groups ranging from “none” (∼30% of genes) to “high” (∼20% of genes) using the average normalised expression values from MEF-1 and MEF-2 RNA-seq datasets (fig. S4). We observed that intermediately methylated CpGs (UI, I, and MI) are more likely to be located within promoters and bodies of genes that are not expressed, or expressed at low levels, whereas U and M CpGs are more likely to be located within highly expressed genes (Fig. 1G and fig. S5, A and B). Moreover, we found that CpGs classified as intermediately methylated (UI, I, and MI) are more likely to be intergenic compared to U and M CpGs (Fig. 1H and fig. S5C). These results reveal a relationship between transcriptional activity and the stability of DNA methylation across the samples.

Methylation levels differ between promoters, which are typically hypomethylated, and gene bodies, which are frequently hypermethylated (*15*). Consistent with this, at highly expressed genes, we found that DNA is hypomethylated at the promoter and first exon, and hypermethylated at the rest of exons and introns (Fig. 2A, fig. S6A, and table S3). The methylation dynamics of highly expressed genes are matched with a high fidelity score throughout the entire gene (Fig. 2B and fig. S6B). This pattern is in sharp contrast to what is observed in genes with no or low expression. In this case, the entire genic region, including promoters and gene bodies alike, is enriched for intermediately methylated CpGs with low methylation fidelity.

**Fig. 2.**
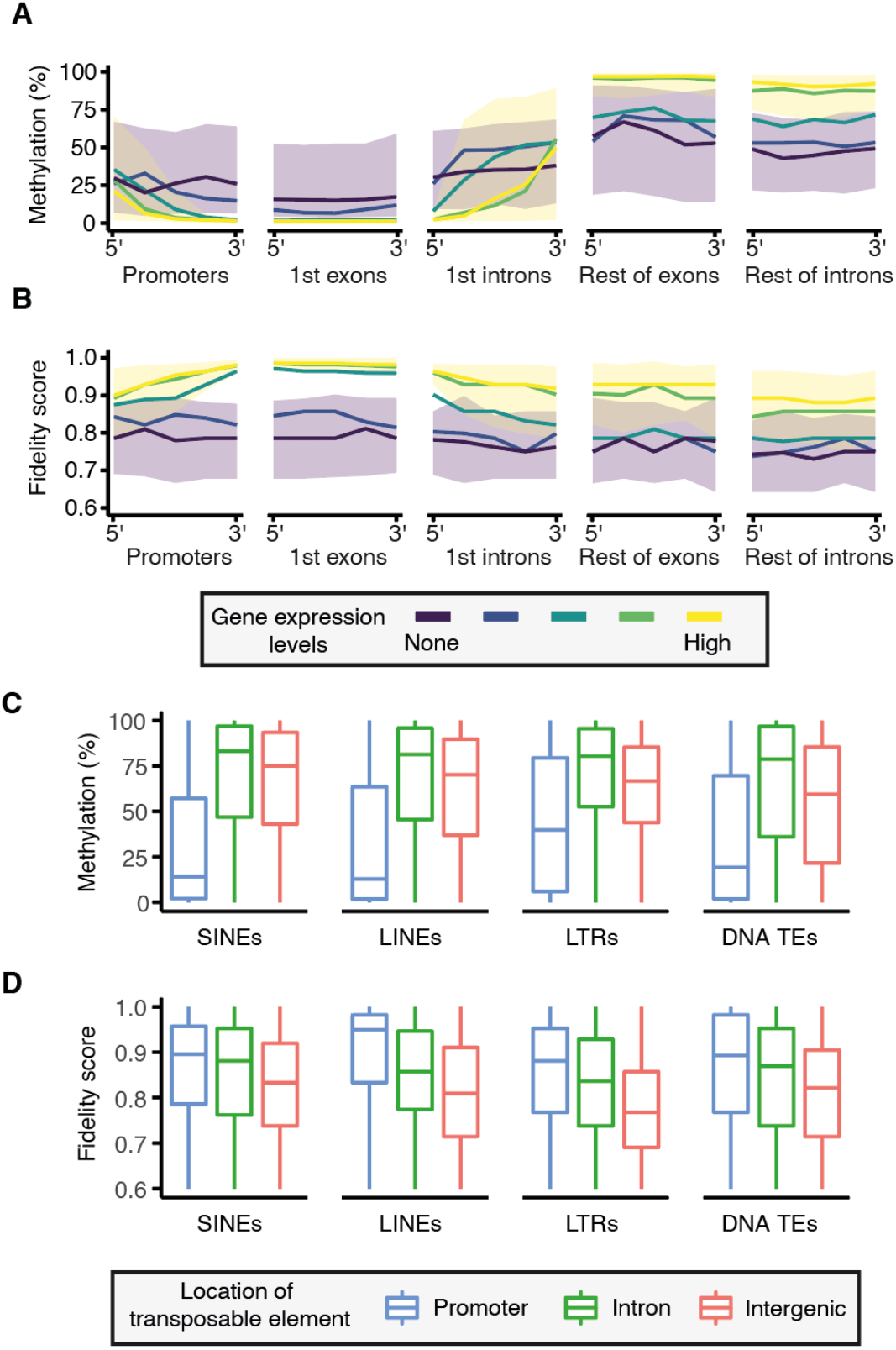
Unfaithful intermediate methylation associates with the lack of transcription. (A) Methylation levels and (B) fidelity score characterised along the following regions of protein-coding genes: promoters, 1st exons, 1st introns, the rest of exons, and the rest of introns. Each region is split into five tiles at which median (lines) or interquartile range (ribbons) of methylation or fidelity score is shown. Only regions covered by at least 3 CpGs are considered; single-exon genes are excluded. Gene expression levels are represented by colours ranging from purple (no expression) to yellow (high expression). (C) Boxplots showing methylation and (D) fidelity score of transposable elements (SINEs, LINEs, LTRs, and DNA transposons) that exist in promoter (blue), intronic (green), or intergenic regions (orange) of the genome. Methylation levels and fidelity score are calculated as the mean across CpGs in each TE.

This suggests that either transcriptional activity defines the patterns of faithful hypo- and hypermethylation throughout a gene, or that a faithful methylation state contributes to expression. Furthermore, it indicates that the absence of transcription may be permissive for the presence of unfaithful intermediate methylation levels.

Transposable elements (TEs) are mobile genetic units that can be transcriptionally repressed by DNA methylation (*16, 17*). However, we found that many CpGs within TEs are intermediately methylated (Fig. 2C). Therefore, we asked whether intermediate methylation associates with either the age and/or has a particular distribution within TE sequences. For example, recently integrated TEs are more likely to retain transcriptional potential and therefore may be preferentially targeted by DNA methylation (*17*). We assessed the relationship between methylation status of a CpG and the evolutionary age of the TE it overlaps (where age is measured as the percent divergence of the individual element from the TE consensus sequence) (fig. S7). Globally, TE sequence divergence does not correlate with DNA methylation level nor fidelity (fig. S7B and C). Intermediate methylation of CpGs within TEs is thus unlikely to be related to the age of the element. Moreover, we could not find an association between intermediate methylation levels and the position of a CpG within a TE. Using SINEs as a tractable model, we observed that methylation levels are similar irrespective of the CpG position along the element (fig. S7D), even though like genes, SINEs are structured and have their own promoter regions (*18*). Taken together, these results indicate that intermediate CpG methylation within TEs is not dependent on TE sequence divergence and is equally distributed along TE sequences, at least in SINEs.

Besides being frequently found in intergenic regions, TEs can also exist in genic regions such as promoters and introns, but rarely in exons (fig. S8) (*19, 20*). Given that promoters and gene bodies generally have distinct patterns of methylation, we tested whether intermediate methylation levels at TEs can be determined by the genomic features of the TE insertion site. We observed that TEs in promoters are generally hypomethylated and TEs in introns tend to be hypermethylated (Fig. 2C). TEs in intergenic regions tend to be less methylated than those in introns, but higher than those in promoters. Indeed, intermediate methylation of CpGs within TEs is more likely to occur within intergenic regions, which is consistent with our observation that CpGs in intergenic TEs have lower fidelity scores compared to those in promoters and introns (Fig. 2D). Therefore, the lack of transcriptional activity at the insertion locus is strongly associated with intermediate methylation within TEs, and in turn its fidelity.

The fidelity score allowed us to determine that intermediate methylation states are unfaithfully propagated between cell divisions. However, this metric does not include the parental line methylation state, and therefore cannot be used to determine how states may be transmitted between the parental and clonal cell lines. Conceptually, we propose two ways by which DNA methylation at a given CpG is transmitted through cellular divisions: faithful and stochastic (Fig. 3A). A faithful process would result in the clonal lines being enriched for 0%, 50%, and 100% methylation states in proportions that recapitulate the parental methylation state. Whereas a stochastic process would result in the clonal lines being enriched for a distribution centred around the parental average methylation level. With low (0-10%) or high (90-100%) parental methylation states, the clonal lines tend to recapitulate the parental methylation level, better fitting a faithful process (Fig. 3B and fig. S9A). However, for the “low intermediate” (10-40%), “intermediate” (40-60%), and “high intermediate” (60-90%) parental methylation states, the clonal lines display modal and skewed distributions that are not reflective of strictly faithful methylation inheritance.

**Fig. 3.**
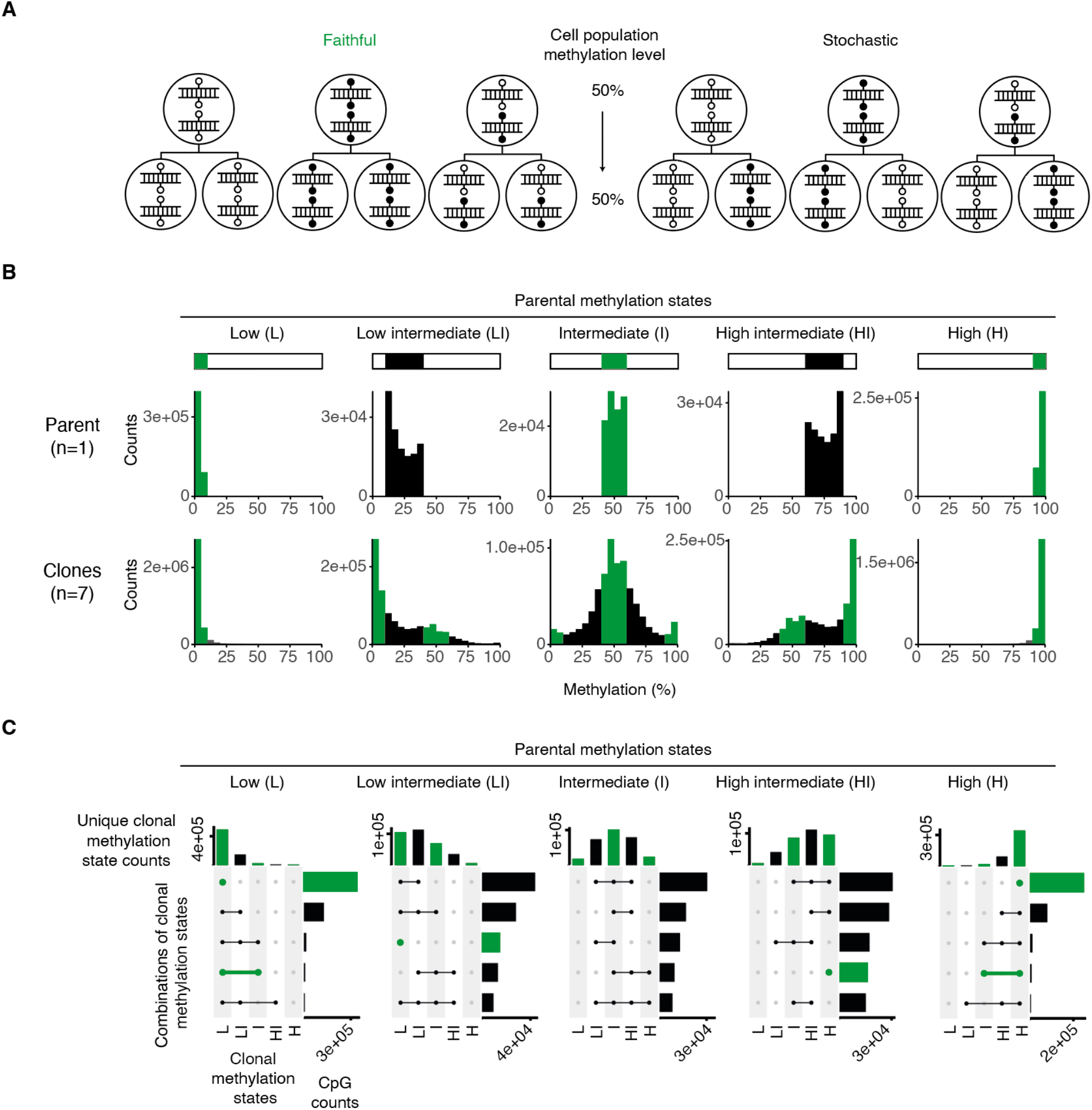
Intermediately methylated CpGs are prone to probabilistic inheritance between cell divisions. (A) Two ways by which DNA methylation at a given CpG site can be transmitted between cell divisions: faithful and stochastic. Each large circle represents a cell - the top row represents the parental cells, and the bottom row represents the daughter cells that arise from cell division. Symmetric methylation at a single CpG site is illustrated as a small filled-in circle on either one or both alleles, whereas absence of methylation is illustrated as a small empty circle. (B) Clonal line methylation distributions from different parental line methylation states for the MEF-1 cell lines. (C) UpSet plots of clonal line methylation states per CpG from different parental line methylation states for the MEF-1 cell lines. In each panel, the top bar plots show the number of unique clonal methylation states represented per CpG. The horizontal bar plots show the CpG counts that exhibit a certain combination of clonal methylation states. Only the five most representative clonal methylation state combinations are shown. Green bars represent cases of potential faithful methylation inheritance because this kind of methylation inheritance will only result in 0%, 50%, or 100% methylation states in the clonal lines. Methylation states are defined quantitatively as the following: Low = 0-10%, Low intermediate = 10-40%, Intermediate = 40-60%, High intermediate = 60-90%, High = 90-100% methylation.

To better visualise these clonal methylation dynamics, we used UpSet plots of the clonal methylation data split by the parental methylation state (Fig. 3C and fig. S9B). With respect to an initial parental methylation state, the top bar plots depict the frequency of distinct states observed amongst the clonal lines for a given CpG. For any given parental state, the most frequently observed state per clonal line CpG is the same as the parental one. The side plots reveal the most common combinations of states observed amongst the derived lines. Unsurprisingly, low (0-10%) and high (90-100%) parental methylation states result most commonly in the same low or high methylation states in the clonal lines, respectively. However, any intermediate parental methylation state (10-40%, 40-60%, 60-90%) most commonly results in a combination of two or three states in the clonal lines, which includes the original state. Hence the heritability of an intermediate state is neither purely faithful nor stochastic. Rather, this suggests a probabilistic inheritance of intermediate methylation states, which allows for some heritability of the cell population methylation level between the parental and clonal lines.

To determine what epigenomic features could be associated with the differential inheritance of methylation states, we calculated fold enrichment for various histone tail modifications that overlap with clonal or probabilistically methylated CpGs (fig. S10A,B & table S4; see methods). We find that histone tail modifications that are generally associated with transcriptional repression (H3K27me3 and H3K9me3 (*21, 22*)) are enriched at the probabilistic intermediately methylated CpGs (fig. S10C). Faithfully methylated CpGs are enriched for histone tail modifications that are associated with transcriptional activation at both hypomethylated promoters (H3K27ac, H3K9ac, H3K4me3 (*23-25*)) and hypermethylated gene bodies (H3K36me3 (*26*)). These findings are in line with our finding noted above that unfaithful methylation is associated with intergenic regions and unexpressed genes (Fig. 2).

The existence of probabilistic unfaithful methylation inheritance suggests that there is continuous loss and gain of methylation at intermediately methylated CpG sites. To test whether DNMT3A/3B are mechanistically responsible for the probabilistic gain of methylation, we generated four MEF lines from *Dnmt3a*^*flox/flox*^*3b*^*flox/flox*^ mice and used recombinant TAT-CRE protein to induce the double knockout (DKO) *in vitro* (Fig. 4A and fig. S11). We performed tcBS-seq with the DKO and control lines, which allowed us to determine the methylation level for ∼2M CpGs (9% of CpGs in the mouse genome). As shown previously (*27*), we found that methylation levels are globally unchanged between the control and *Dnmt3a/3b* deficient MEFs (Fig. 4B). To determine whether methylation levels vary at individual CpG sites, we calculated the difference in methylation between the DKO and controls (Fig. 4C and D). There was no consistent depletion amongst the different methylation states (U, I, and M). Indeed, between the DKO and control cell lines, fewer than 3000 CpGs (∼0.14%) show significantly different levels of methylation (q < 0.01; Fisher’s exact test), with about half increasing and the other half decreasing in methylation level. Additionally, an ANOVA analysis revealed that the independent control cell lines (A, B, C, and D) contribute substantially more to the variance in methylation (F=60.7, p < 2e-16) than the knockout condition (F=1.6, p=0.20). These results show that the deposition of neither probabilistic nor faithful methylation is dependent on DNMT3A/3B in somatic cells, and instead suggest that DNMT1 is the sole methyltransferase involved in both processes.

**Fig. 4.**
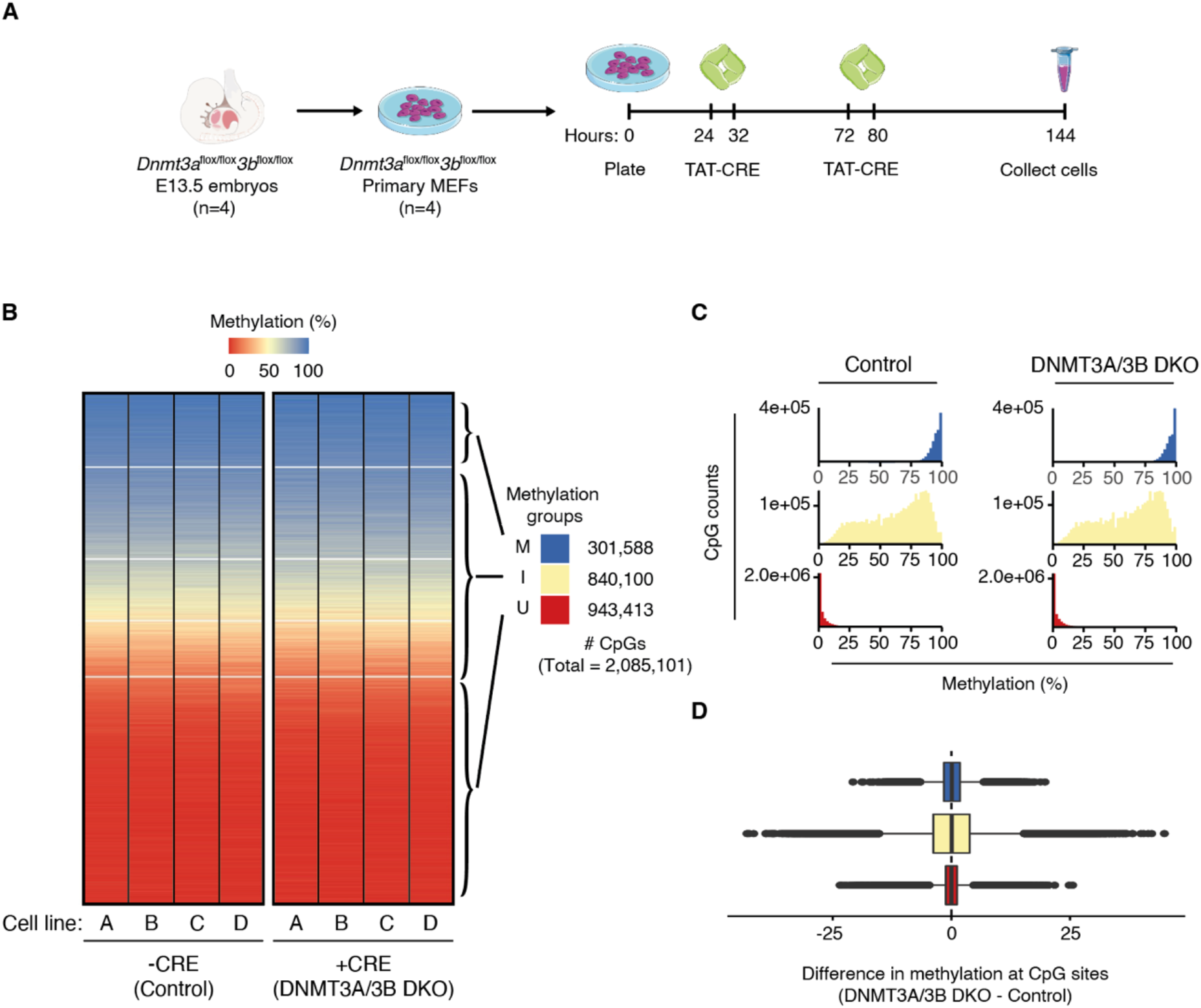
Intermediate methylation does not arise due to *DNMT3a/3b de novo* activity. (A) Primary MEFs were established from *Dnmt3a*^*flox/flox*^*3b*^*flox/flox*^ E13.5 embryos (n=4). The cells were then plated and 24 hours later, treated with recombinant TAT-CRE protein for 8 hours. 48 hours after the initial treatment, the cells were treated with TAT-CRE again for 8 hours. 72 hours after the second TAT-CRE treatment, the cells were collected for DNA/RNA extraction. We generated tcBS-seq libraries for four individual control MEF lines and four matched TAT-CRE treated MEF lines. (B) Heatmaps of 2,085,101 CpGs sorted by median methylation (%) within k-means clusters (k=5) for untreated (control) and TAT-CRE treated (DNMT3A/3B) primary MEF cell lines A, B, C, and D. Methylation groups are classified using the k-means clusters as shown. U (red) = consistently unmethylated across all the cell lines. I (off-white) = potential to be intermediately methylated. M (blue) = consistently methylated across all the cell lines. (C) Methylation percentage distributions of the methylation groups for both the control and DNMT3A/3B DKO cell lines. (D) Boxplot showing the difference in methylation at CpG sites between DNMT3A/3B DKO and control cell lines. Differences per CpG were calculated as the difference of average methylation across the four DNMT3A/3B DKO cell lines and the four control cell lines at a given CpG site. U (red); I (off-white); M (blue).

Based on the premise of its faithful clonal inheritance between cell divisions and its potential to influence transcription, DNA methylation is frequently used as a biomarker for epigenetic ageing and in clinical and epidemiological studies (*28-32*). Our results show that intermediately methylated CpGs are unfaithfully and probabilistically propagated during cell divisions with implications for the use of methylation as a biomarker. We find that these intermediately methylated loci are generally associated with a lack of gene expression, meaning that any functional interpretations are also likely to be unreliable. Due to the observed relationship between gene expression and methylation fidelity, it is important to consider that intermediate methylation states may vary between cell types in accordance with transcriptome-wide fluctuations. The fact that cell populations derived from single cells can exhibit intermediate methylation states after expansion indicates that some loci have the capacity to constantly gain and lose DNA methylation through cell divisions; we suggest that the gain is likely due to the recently uncovered *de novo* function of DNMT1 (*33-36*), while the mechanism for the loss is still unclear.

Differential occupancy of transcription factors could influence the dynamics of methylation deposition at intermediately methylated regions, giving rise to probabilistic methylation states between cell divisions (*3*). Although we could not assign functional features to the intermediately methylated sites we identify, such functionality cannot be ruled out. Our findings challenge the long-standing assumption in the epigenetics field that DNA methylation is a mitotically inherited modification in somatic lineages by revealing that at intermediately methylated sites, methylation levels are probabilistically, not clonally, maintained within a cell population.

## Acknowledgements

We thank members of the Ferguson-Smith group for technical support and useful discussion. We thank Tessa Bertozzi and Ben Simons for critical reading of the manuscript. We thank Mitsuteru Ito for technical assistance. We thank Rahia Mashoodh for support with statistical analysis. We are grateful to Yoach Rais for providing us with TAT-CRE recombinase. We thank Ben Harvey and Joe Gaughan from Agilent for technical support and valuable input.

## Author contributions

A.D.H., F.K.T., and A.C.F.-S conceived and designed the project. A.D.H. performed the experimental work and bioinformatics analyses.N.J.K. contributed to the methylation data processing and analyses. D.G. contributed to the analysis of transposable elements. H.T. contributed to the RNA-sequencing processing and analysis. N.T. contributed to the inducible knockout mouse work. A.D.H., F.K.T., and A.C.F.-S drafted the manuscript. All authors commented on the manuscript.

## Materials and Methods

### Mouse lines

Mouse work was conducted under project licenses from the UK government Home Office (project license numbers: PC9886123, PC213320E, and PP8193772). Mice were housed in a temperature and humidity-controlled room under 12 hr light / 12 hr dark cycles and fed a standard chow diet *ad libitum*. Post-implantation embryos (E13.5 for mouse embryonic fibroblasts) and blastocysts (E3.5 for mouse embryonic stem cells) were collected by natural mating, and the plugged date of conception was considered E0.5. *Dnmt3a*^*flox/flox*^*3b*^*flox/flox*^ mice (*37*) were obtained from RIKEN BioResource Research Center (BRC) and maintained on a C57BL/6 mouse background.

### Cell line generation and culture

Mouse embryonic fibroblasts (MEFs) were established from E13.5 C57BL/6J mouse embryos (*38*). MEFs were grown at 37°C and 5% CO_2_ in high glucose DMEM GlutaMAX™ (ThermoFisher, cat no. 31966021) supplemented with 10% FBS (Gibco, 11550356) and 1% penicillin/streptomycin solution and passaged with trypsin/EDTA. MEFs were immortalised by serial passaging the cells through crisis phase (*39*). Isolation of single immortalised MEF cells was performed by flow cytometry (MoFlo Astrios Cell Sorter) to 96-well plates. The isolated single cells were grown into clonal cell lines by successive passaging from 96-well, 48-well, 24-well, 12-well, and 6-well plates to 10cm dishes where they were grown to 75% confluency to yield 1-2 million cells per clonal line for DNA/RNA extraction. The parental lines, from which the clonal lines were derived, were also grown to 75% confluency on 10cm dishes to yield 1-2 million cells for DNA/RNA extraction.

### DNA/RNA extraction

DNA/RNA extraction was performed with the Qiagen AllPrep DNA/RNA Mini Kit (cat no. 80204).

### Target capture bisulfite sequencing (tcBS-seq)

Libraries were generated using the SureSelectXT Methyl-seq Library Preparation Kit with the following specifications. 1μg of DNA was sonicated to an average size of 200bp with the Covaris E220 (Duty Factor = 30%, PIP = 100, Cycles per Burst = 1000, Treatment Time = 95, Bath Temperature = 7°C, with intensifier fitted) using 50μl microTUBEs (cat no. 520166) and 24-place rack (cat no. 500308). Following end repair, dA tailing, and adaptor ligation, libraries were hybridised to single-stranded RNA probes homologous to 297,000 regions in the mouse genome (SureSelectXT Methyl-Seq Capture Probes, cat no. 931052). After purification, the libraries were bisulphite converted using Zymo Research’s Methylation-Gold Kit (cat no. D5005) and PCR amplified for 8 cycles. The PCR amplified bisulphite-treated libraries were then purified, indexed by PCR amplification (6 cycles), and purified again. All purification steps took place using Ampure XP beads (Beckman Coulter, cat no. A63881). The final libraries were then quality checked and quantified for multiplexing using the Bioanalyzer High-Sensitivity DNA kit (Agilent, 5067-4626) and Qubit dsDNA HS Assay kit (Thermo Scientific, Q32854). The multiplexed libraries were sequenced as 150 bp paired-end reads on the Illumina Novaseq 6000 SP. The resulting tcBS-sequencing data was trimmed by Trim Galore (v0.6.0) and aligned to mm10 using Bismark (*40*). Reads with map quality score less than 10 were excluded and were further filtered by M-bias filtering (*41*): for MEF-1 and MEF-2 data, we excluded the first two bp on both paired reads and the last bp on Read 2; for ESC-1 and ESC-2, we excluded the first six bp on Read 1, the first seven bp on Read 2, and the last bp on Read 2. Methylation data was extracted using bismark_methylation_extractor (*40*) and analysed in R (v3.6.1).

### Filtering and thresholding tcBS-seq methylation data

The SureSelectXT tcBS-seq system uses RNA probes homologous to 300,000 genomic regions, which represent 109 Mbases of the 2.7 Gbase mouse genome, and about three million CpGs of the total 20 million CpGs. We use read coverage across the individual sequencing libraries to look for enrichment of reads on individual chromosomes to determine coverage-based karyotypes for our immortalised MEF cell lines and find that chromosomes 12, 18, and 19 exhibit aneuploidy in at least one of the cell lines (fig. S1, A and B). For subsequent analyses, we remove the CpGs on the aneuploid chromosomes, as well as those on the X chromosome due to the sex difference between the two parental MEF lines (fig. S1C). We used the following primers to amplify the SRY gene on the Y chromosome:

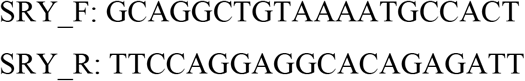

After thresholding for CpGs with greater than or equal to 10x coverage in all 16 sequencing libraries, for subsequent analyses, we retain 1.2 million CpGs (or ∼5% of CpGs in the mouse genome) with a median coverage of 32 reads per CpG per dataset (fig. S1D). To assess whether the tcBS-seq CpGs are representative of CpGs genome-wide, we examine the distribution of CpGs amongst genomic annotations and compare the methylation data to similar whole-genome bisulfite methylation datasets. First, we calculate the proportion of various genomic annotations covered by at least one tcBS-seq CpG. We find that these CpGs are found within 27.3% of promoters, 13.8% of exons, and 22.9% of introns in the genome, whereas they only overlap with ∼1-2% of transposable elements (TEs) (table S1). This informs us that tcBS-seq enriches for genic regions and is depleted for TEs. Next, we compare the methylation profiles of all MEF-1 (n=8) and MEF-2 (n=8) tcBS-seq datasets to other relevant methylation data (table S2). To do this, we utilised publicly available WGBS datasets from both *in vitro* (primary and immortalised MEFs) (*42, 43*) and *in vivo* contexts (E13.5 and E14.5 embryonic limb bud3) (*44*). When filtered for the same CpGs covered in our dataset, we observe that all the datasets have similar methylation distributions with enrichment for hypo- and hypermethylation, and depletion of intermediate states (fig. S2A). We observe that the MEF-1, MEF-2, and publicly available primary MEF datasets, are globally reduced in methylation compared to immortalised MEFs and E13.5/E14.5 embryonic limb bud (fig. S2B). However, when considering the entire genome, the tcBS-seq MEF-1 and MEF-2 datasets are depleted in global methylation levels compared to the WGBS datasets. This suggests that MEF-1 and MEF-2 methylation profiles are more like those of primary MEFs as opposed to immortalised MEFs, and more importantly, that tcBS-seq enriches for hypomethylated regions of the genome.

Different kinds of regions in the genome exhibit distinctive methylation patterns. For example, genomic imprints are allelically methylated and exhibit 50% methylation levels, while TEs are hypermethylated compared to the background methylation level of the genome. Additionally, gene promoters are generally hypomethylated, whereas exons and introns tend to be hypermethylated. We compare methylation profiles of the relevant public datasets at imprints, gene bodies, and SINEs (a major family of TEs), to validate that, despite being depleted for methylated regions, the MEF-1 and MEF-2 datasets show the expected distinctive methylation distributions. We filter publicly available methylation datasets for the tcBS-seq covered CpGs and find that all the datasets have similar distributions of methylation at imprints, across gene bodies, and at SINEs (fig. S2C-E). This suggests that despite the underrepresentation of TEs and methylated regions in the tcBS-seq, there is enough methylation data that is representative and typical of the somatic mouse methylome to address questions regarding methylation inheritance between cell divisions.

### Characterising the clonal methylation data

Here we explain the calculations for the data visualised by the heatmaps of Fig. 1D-F. CpG density was calculated as the number of CpGs within 100bp of the focal CpG, with an upper limit of 30 CpGs. Fidelity score was calculated as the number of clonal lines that exhibit [0-10], [40-60], or [90-100] % methylation, divided by 14 (the total number of clonal lines). Neighbour similarity score was calculated as the number of clonal lines in which the closest CpG to a focal CpG is within 10% methylation, divided by 14 (the total number of clonal lines).

### Classifying CpGs by methylation

Throughout this manuscript we classify CpGs in five different ways for subsequent analyses: 1) As k-mean clusters, 2) combined k-means methylation groups, 3) methylation state bins, 4) probabilistic and faithful, and 5) combined k-means methylation groups for the conditional DNMT3A/3B knockout experiment.

1. Figure 1C shows the 7 k-means clusters of methylation data arranged by median methylation.
2. Figure S3A and B show how we combine the k-means clusters to unmethylated (U), unmethylated and intermediately methylated (UI), intermediately methylated (I), methylated and intermediately (MI), and methylated (M).
3. For Figure 3B, C and Figure S9A and B, we characterise CpG methylation states as low [0, 10), low intermediate [10, 40), intermediate [40, 60), high intermediate [60, 90), and high (90, 100].
4. For Figure S10, we define CpGs as probabilistic if a clonal line exhibits (10-40] or (60-90] % methylation amongst both the MEF-1 and MEF-2 clonal lines. CpGs are defined as faithful if all clonal lines exhibit [0-10], (40-60], or (90-100] % methylation.
5. For Figure 4B and C show how we combine the k-means clusters to unmethylated (U), intermediately methylated (I), and methylated (M).

### Analysing methylation at transposable elements

To estimate the proportion of TEs for which methylation data is available (table S1), CpG coordinates were compared with the RepeatMasker v4.1.1 annotation for mm10 (obtained from the UCSC genome browser). Data on transposon sequence divergence from consensus was taken directly from RepeatMasker output and plotted for each transposon class (i.e., DNA transposons, LTR retrotransposons, LINEs, and SINEs). Consensus alignment positions (1-200 from 5’-ends) for genomic copies of SINEs were similarly extracted from the mm10 RepeatMasker output.

### Total RNA sequencing

Libraries were generated using the NEBNext® rRNA Depletion Kit (NEB, cat no. E6310) and UltraTM II Directional RNA Library Prep Kit for Illumina® (NEB, cat no. E7760) with the following specifications. RNA Integrity Number (RIN) was determined using the Agilent RNA 6000 Pico Kit (cat no. 5067-1513) on the Agilent 2100 Bioanalyzer and its associated software. 1μg of RNA was hybridised to probes for rRNA depletion, treated with RNase H and DNase I, and purified before being fragmented at 94°C with incubation times ranging from 8 to 15 minutes depending on the RIN. The fragmented RNA was then reverse-transcribed to cDNA in two steps and purified. Following end repair and adapter ligation, the cDNA libraries were purified, PCR enriched for 9 cycles, and purified again. All purification steps took place using Ampure XP beads (Beckman Coulter, cat no. A63881). The final libraries were quality checked and quantified for multiplexing using the Bioanalyzer High-Sensitivity DNA kit (Agilent, 5067-4626), Qubit dsDNA HS Assay kit (Thermo Scientific, Q32854), and the KAPA Library Quantification Kit optimised for Roche® LightCycler 480 (Roche, 07960298001). The multiplexed libraries were sequenced as 150 bp paired-end reads on the Illumina Novaseq 6000 S1. The resulting RNA-sequencing data was trimmed by Trim Galore (v0.6.0), aligned (mm10) and quantified using Salmon (v1.5.2) (*45*) using the mm10 reference annotation (Ensembl release 102, Nov 2020). The transcript quantification was processed using the R/Bioconductor package DESeq2 (v1.24.0) (*46*), to obtain normalised counts using the “regularised log” (rlog) transformation.

### Characterising transcriptomic data

For transcriptomic analyses, only genes annotated as protein-coding by Ensembl (release 102) were considered, and single-exon genes were excluded, for a total of 20,273 genes. To get a single transcript per gene, canonical transcripts were first defined as the most highly expressed transcript from a gene on average across all the MEF parental and clonal total RNA-seq datasets; for genes lacking transcript expression in those datasets, the mm10 “known canonical” transcripts as defined by the UCSC genome browser were used. From these single transcripts, the “first exon” for each gene was determined; the promoter region was defined as 1000bp prior to this first exon. The annotations were classified into the following genic regions: promoters, 1st exons, 1st introns, 2nd exons, 2nd introns, 3rd exons, 3rd introns, rest of exons, rest of introns, last introns, last exons (see table S3). Only genic regions covered by at least three CpGs in the methylation data and greater than 6bp in length were considered. Next, each genic region was divided into five tiles. Transcription quintiles were derived from the normalised expression values for each annotated protein-coding gene averaged across all the MEF clonal and parental total RNA-seq datasets and assigned to the corresponding tiled regions of the genes (figs. S4, S5, and S6 and Fig. 2A and B). CpGs were overlapped with protein-coding transcripts and their promoters (1000bp prior to the TSS of each of the transcripts) and assigned a corresponding transcription quintile (fig. S5B and Fig. 1G). CpGs that did not overlap with a protein-coding transcript or a promoter, were classified as intergenic (fig. S5C and Fig. 1H).

### In vitro knockout of *Dnmt3a/3b*

Primary MEFs were established from *Dnmt3a*^flox/flox^*3b*^flox/flox^ E13.5 embryos (*38*) and grown at 37°C and 5% CO_2_ in high glucose DMEM GlutaMAX™ (ThermoFisher, cat no. 31966021) supplemented with 10% FBS (Gibco, 11550356) and 1% penicillin/streptomycin solution and passaged with trypsin/EDTA. To induce the knockout of DNMT3A/3B, 1×10^6^ cells were plated and the next day washed with PBS and media was replaced with 5ml of DMEM:PBS (1:1) with 50μl TAT-CRE recombinase (2-3μg/μl) and incubated at 37°C for 8 hours. Following the TAT-CRE recombinase treatment, cells were washed three times with PBS and the original growth media was replaced. The TAT-CRE recombinase treatment was carried out again 48 hours following the start of the initial treatment. Cells were collected for protein and DNA/RNA extraction 72 hours after the second TAT-CRE recombinase treatment. Knockout of the genes was confirmed by running PCR reactions using Q5® High-Fidelity DNA Polymerase (NEB, cat no. M0491) and the following primers on a 2.5% agarose gel:

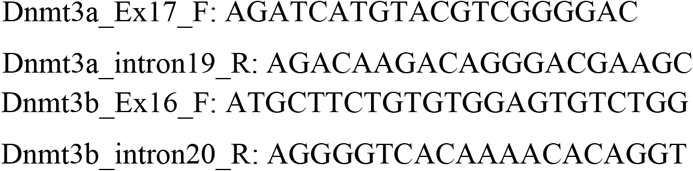

### Western blotting

Flash frozen control and knockout MEFs were thawed on ice and lysed with RIPA buffer (Sigma-Aldrich, cat no. R0278), supplemented with EDTA-free cOmpleteTM Protease Inhibitor Cocktail (Roche), for 20 minutes on ice and centrifuged 14,000g for 10 minutes at 4°C. Supernatant was collected and protein concentrations were determined using Bradford Reagent (Sigma-Aldrich, cat no. B6916). Protein samples were boiled with 4x Laemmli sample buffer (Bio-Rad, cat no. 1610747), with 10% b-mercaptoethanol, at 95°C for 5 minutes. Protein ladder (Thermo Fisher Scientific, cat no. 26619) and equal amounts of protein sample were then loaded onto a 4-20% gradient SDS-PAGE gel (Bio-Rad, cat no. 4561095) and run at 120V before being transferred to a PVDF membrane (Bio-Rad, cat no. 1704156). The membrane was then blocked with 5% skim milk and incubated with the following primary antibodies for 1 hour at room temperature: anti-DNMT3a (1:1000, ab188470) and anti-b-actin (1:1000, ab8227). After three 10 minutes washes with 1xTBST buffer at room temperature, the membrane was incubated with secondary antibody, goat anti-rabbit IgG-HRP (1:3000, Agilent, cat no. P044801-2), for 1 hour at room temperature. The membrane was again washed three times for 10 minutes at room temperature with 1xTBST buffer; signal was detected using Amersham ECL (GE Healthcare, cat no. RPN2232) and imaged on a LI-COR Odyssey® Fc Imaging System.

---

**Fig. S1:**
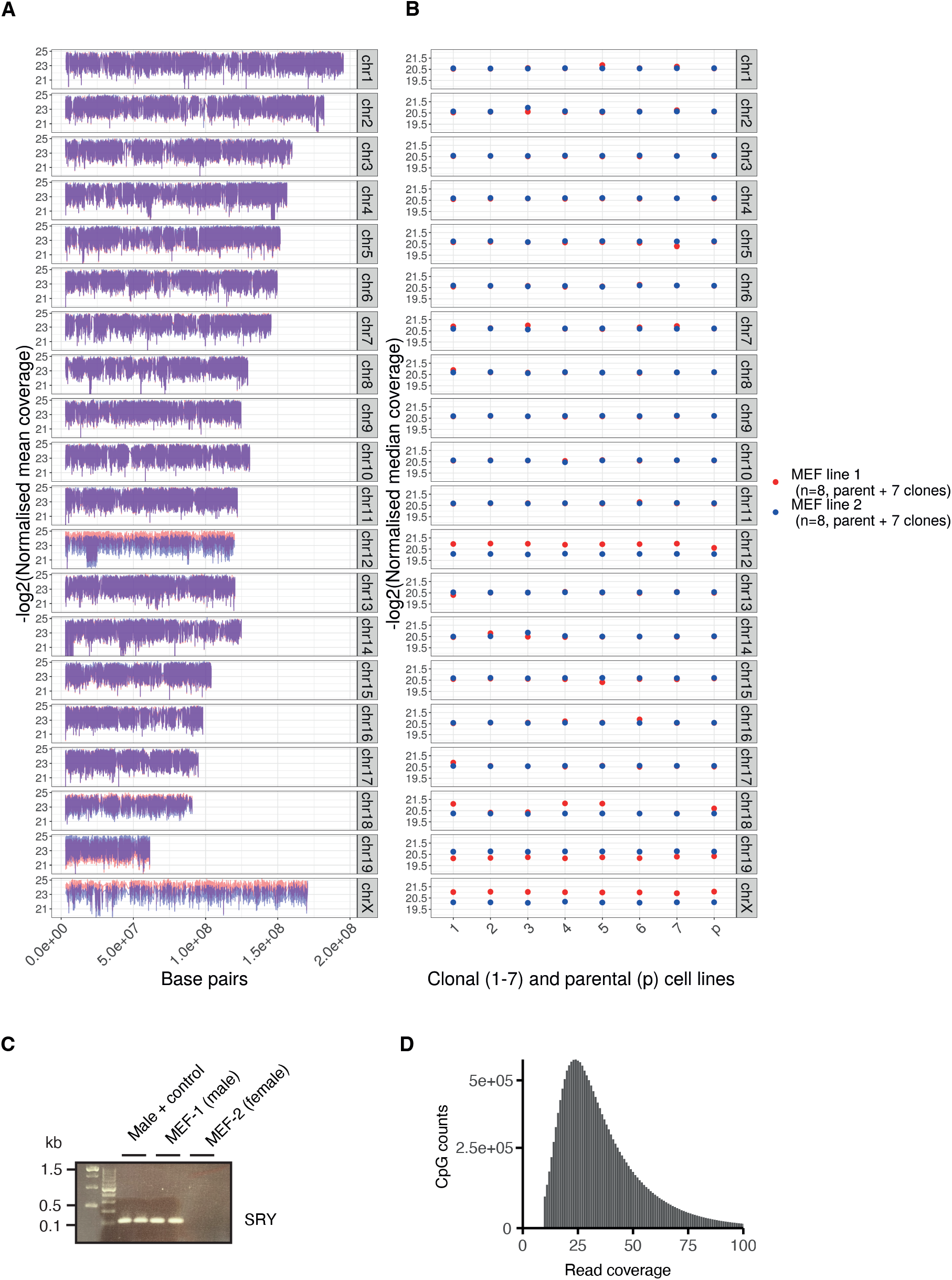
Filtering and thresholding MEF methylation data. A) Mean coverage (log2 normalised) across MEF-1 and MEF-2 as well as (B) the median coverage (log2 normalised) for each of the individual libraries. Decreased coverage across chromosome 12 and 18 for all or some of the MEF-1 lines compared to the MEF-2 lines suggests monosomies for those two chromosomes in some or all the MEF-1 lines. Increased coverage across chromosome 19 for all the MEF-1 lines suggests trisomy 19 for all the MEF-2 lines. For subsequent analyses, the CpGs on chromosomes 12, 18, and 19 were filtered out. Increased coverage on the X chromosome for all the MEF-2 lines suggests that these lines were derived from a female embryo. (C) Sex of the two lines was independently determined by PCR of the SRY gene, which confirmed that MEF-1 was derived from a male embryo and MEF-2 from a female embryo. For subsequent analyses, the CpGs on chromosomes X and Y were filtered out. (D) Thresholding data at greater than 10x coverage in all 16 methylation datasets results in a median coverage of 32 reads per CpG.

**Fig. S2:**
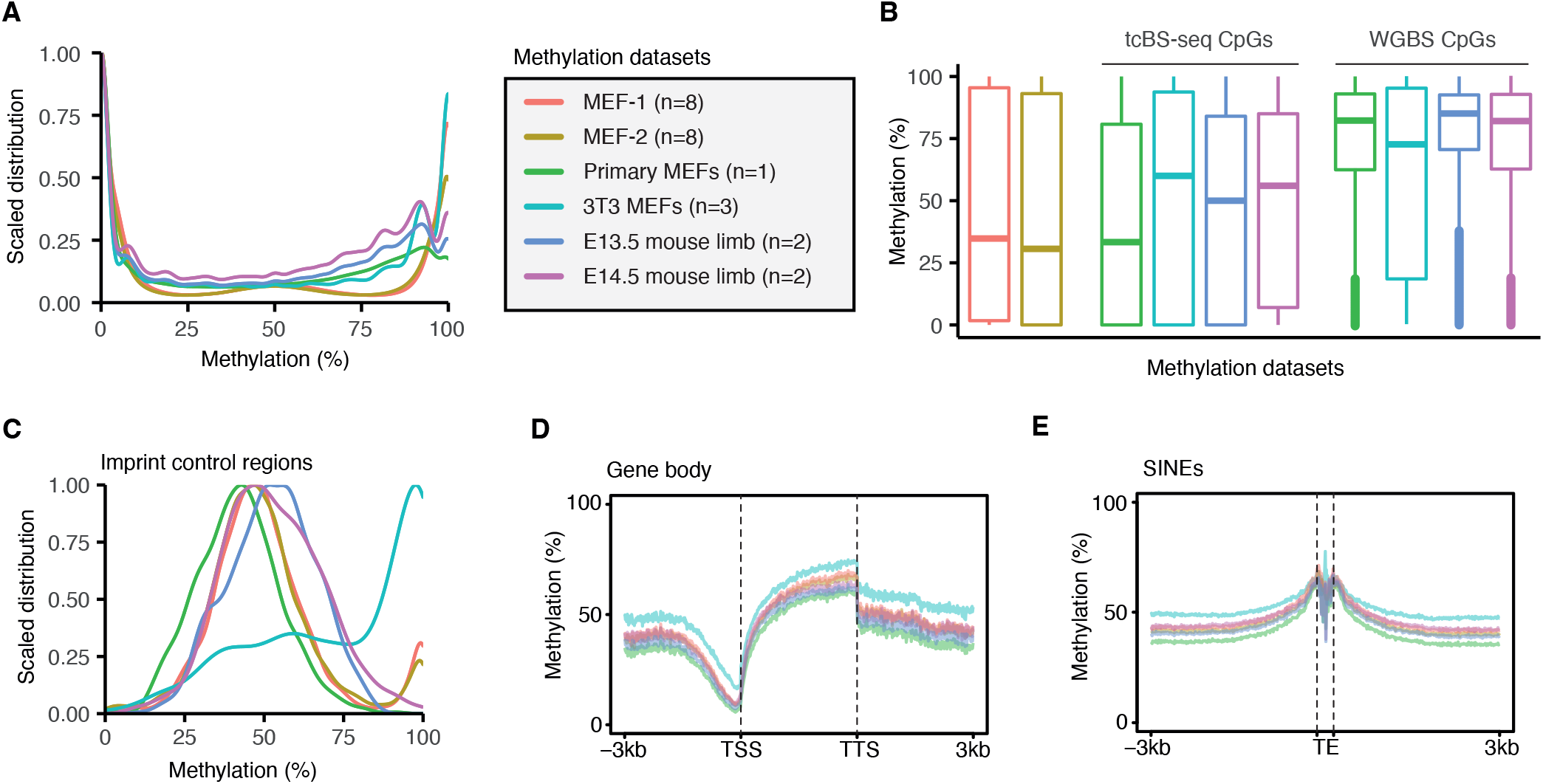
Validation of MEF target capture bisulphite sequencing. A) Density plots to show methylation distribution of the thresholded and filtered tc-BSseq CpGs for MEF-1, MEF-2, and relevant publicly available whole genome bisulphite sequencing (WGBS) datasets (see Table 3.2). All datasets have comparable profiles of methylation for the tcBS-seq CpGs. The accompanying legend applies to all panels of this figure. (B) Boxplots of methylation distribution for MEF-1 and MEF-2 datasets compared to publicly available whole genome bisulphite sequencing (WGBS) datasets that are also filtered for the tcBS-seq CpGs. MEF-1 and MEF-2 have a similar distribution to the primary MEFs when filtered for the tcBS-seq CpGs, but a lower distribution when considering CpGs genome-wide. (C) Methylation distributions of CpGs at imprint control regions are similar across all datasets, aside from the 3T3 immortalised MEFs. (D) Methylation shows a similar pattern of reduced values at TSSs and enrichment across gene bodies in all datasets when filtered for the tcBS-seq CpGs. (E) Methylation is enriched at the 5’ and 3’ edges of SINEs in all datasets when filtered for the tcBS-seq CpGs.

**Fig. S3:**
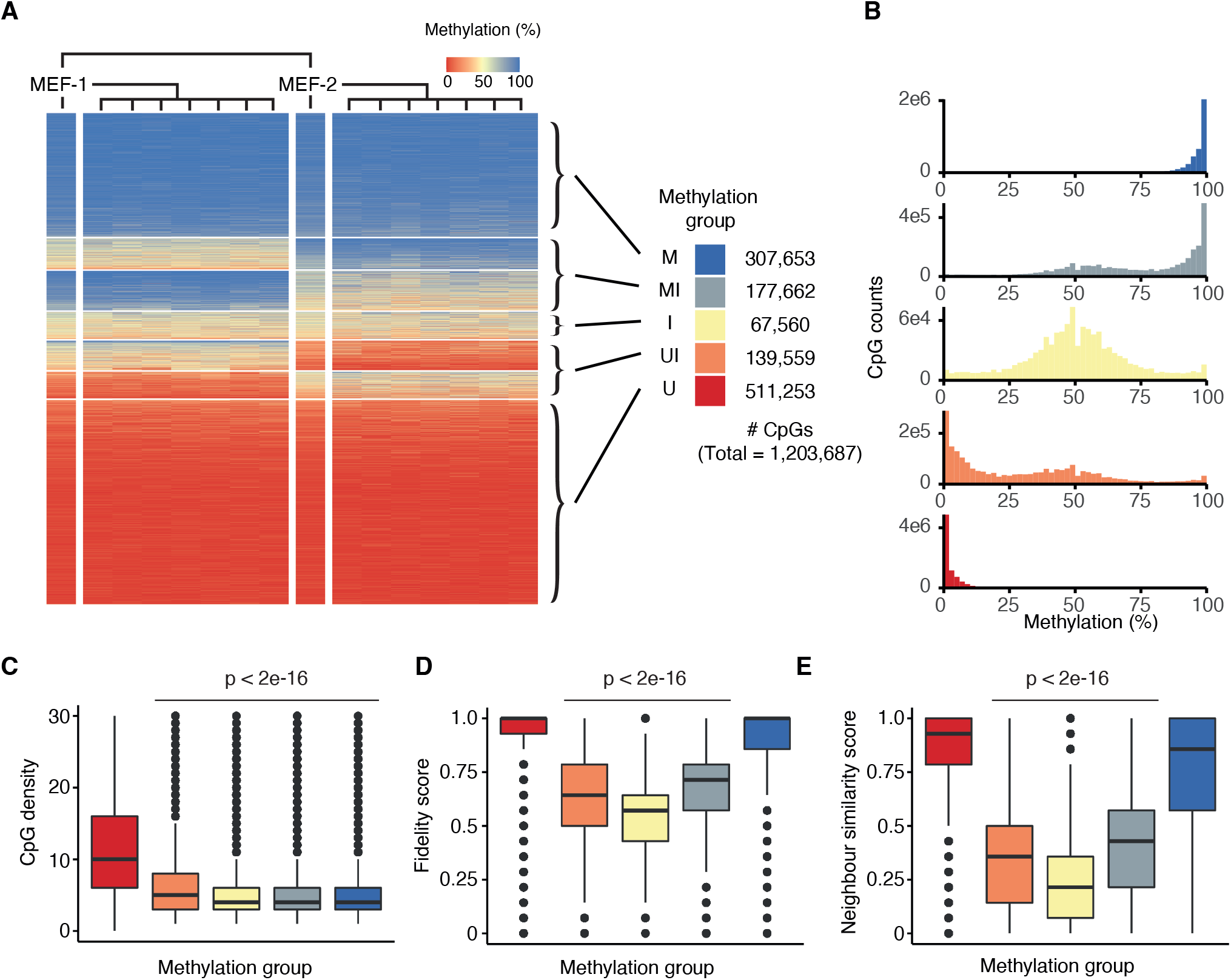
Classifying and evaluating methylation groups in MEFs. A) Classifying five methylation groups for 1,203,687 CpGs from the seven k-means clusters of MEF-1 and MEF-2 parental and clonal line methylation data (as shown in Fig. 1C). U = consistently hypomethylated across all the cell lines. UI = potential to be either hypo- or intermediately methylated. I = intermediately methylated. MI = potential to be either hyper- or intermediately methylated. M = consistently hypermethylated across all the cell lines. (B) Methylation distributions of the methylation groups across all MEF datasets. (C) CpG density per 100bp, (D) Comparing fidelity score, and (E) neighbour similarity score of the different methylation groups. P-values for (C) are from quasi-Poisson regressions comparing values from U, I, MI, and M with U; p-values for (D) and (E) are from Wilcoxon rank-sum tests comparing values from U, I, and MI with U and M.

**Fig. S4:**
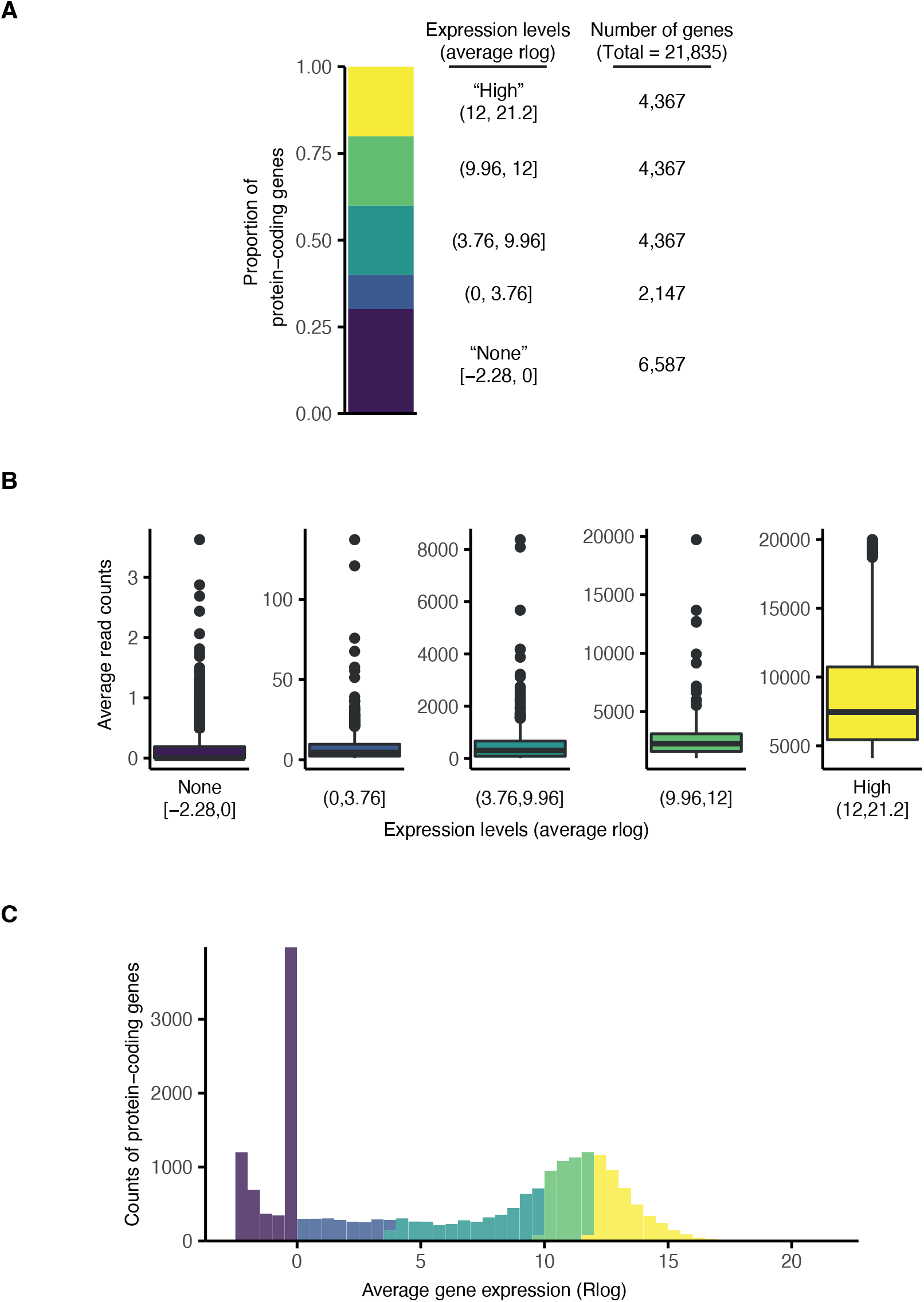
Classifying expression levels in MEFs. (A) Bar plot showing proportions of protein-coding gene expression levels as determined by average rlog expression ranges quantified using all MEF-1 and MEF-2 RNA-seq datasets. The annotation of all 21,835 protein-coding genes in the mouse genome is derived from Ensembl release 102. (B) Boxplots showing average read counts of the different expression levels. (C) Distribution of average gene expression rlog values coloured by gene expression levels.

**Fig. S5:**
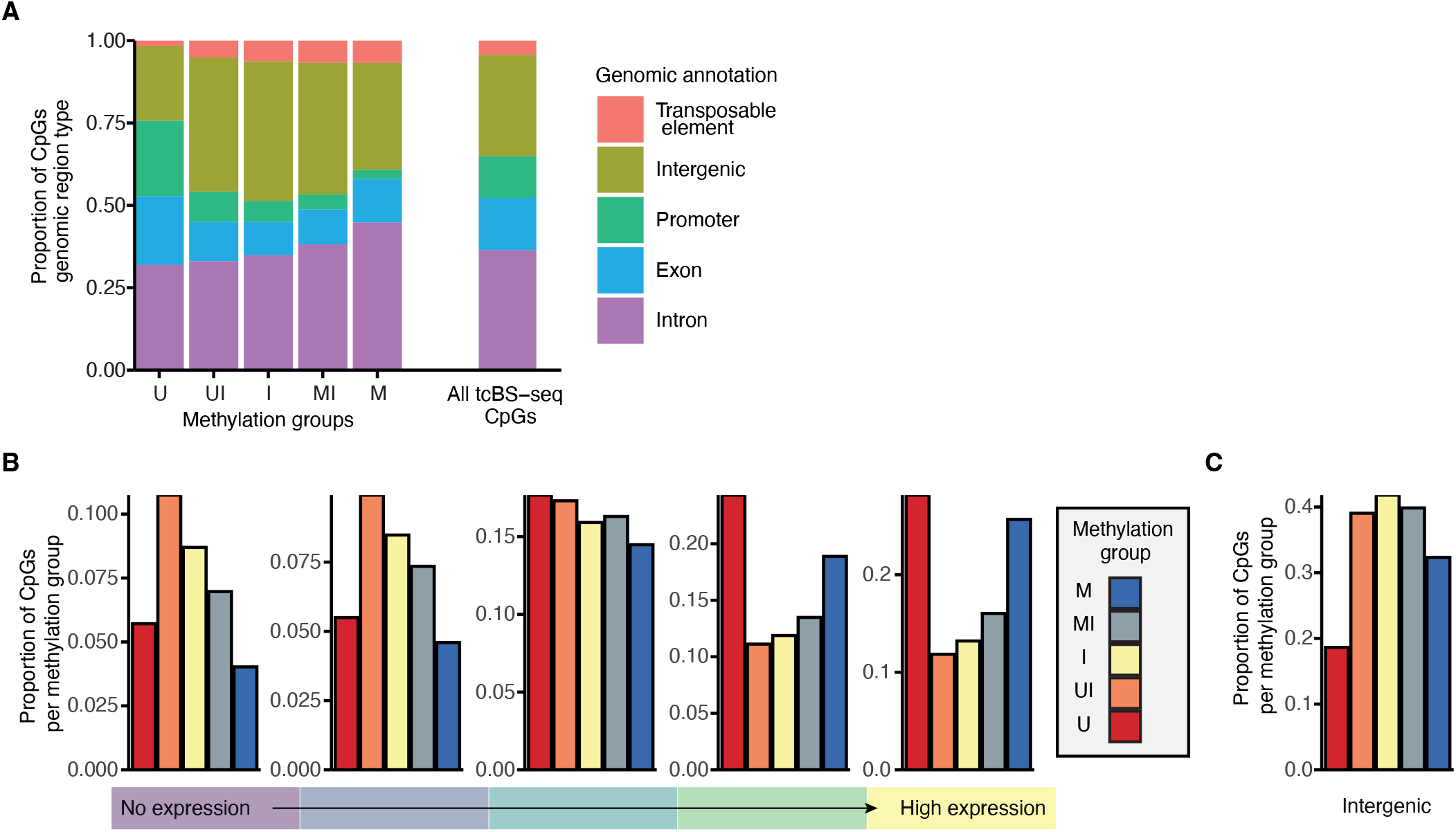
CpGs that exhibit intermediate methylation associate with transcriptional inactivity in MEFs. (A) Relative distribution of different genomic annotations amongst the methylation groups and all CpGs assessed. (B) Proportion of CpGs in methylation groups that overlap with protein-coding genes of varying expression levels. For this overlap, 1000bp before the transcription start site is included at all protein-coding genes to account for CpGs that overlap with promoters. (C) Proportion of CpGs in methylation groups that are intergenic. CpGs that do not overlap with a protein-coding gene or promoter are classified as intergenic. Gene expression levels are represented by colours ranging from purple (no expression) to yellow (high expression).

**Fig. S6:**
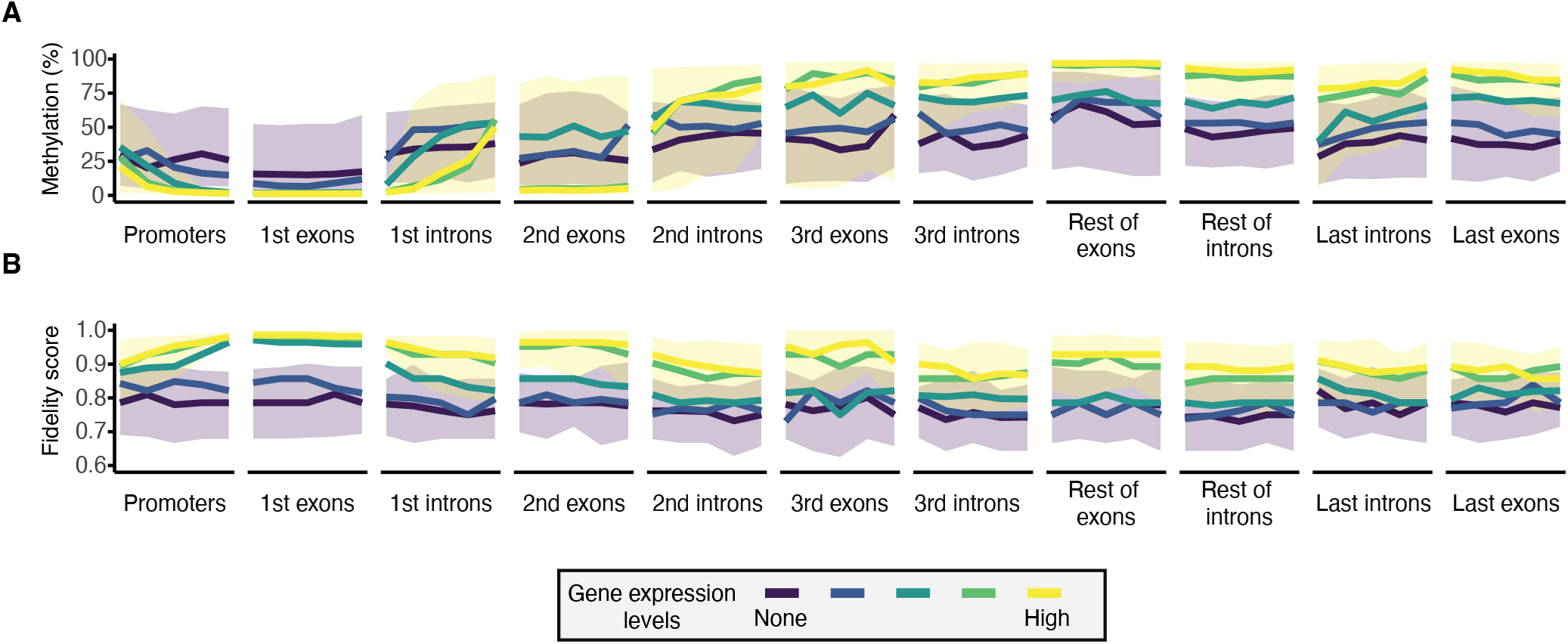
Methylation levels and methylation fidelity at protein-coding genes of varying expression in MEFs. (A) Methylation levels and (B) fidelity score characterised along regions of protein-coding genes. Each genic region is split into five tiles at which median (lines) or interquartile range (ribbons) of methylation or fidelity score is shown. Only genic regions covered by at least 3 CpGs are considered; single-exon genes are excluded. Gene expression levels are represented by colours ranging from purple (no expression) to yellow (high expression).

**Fig. S7:**
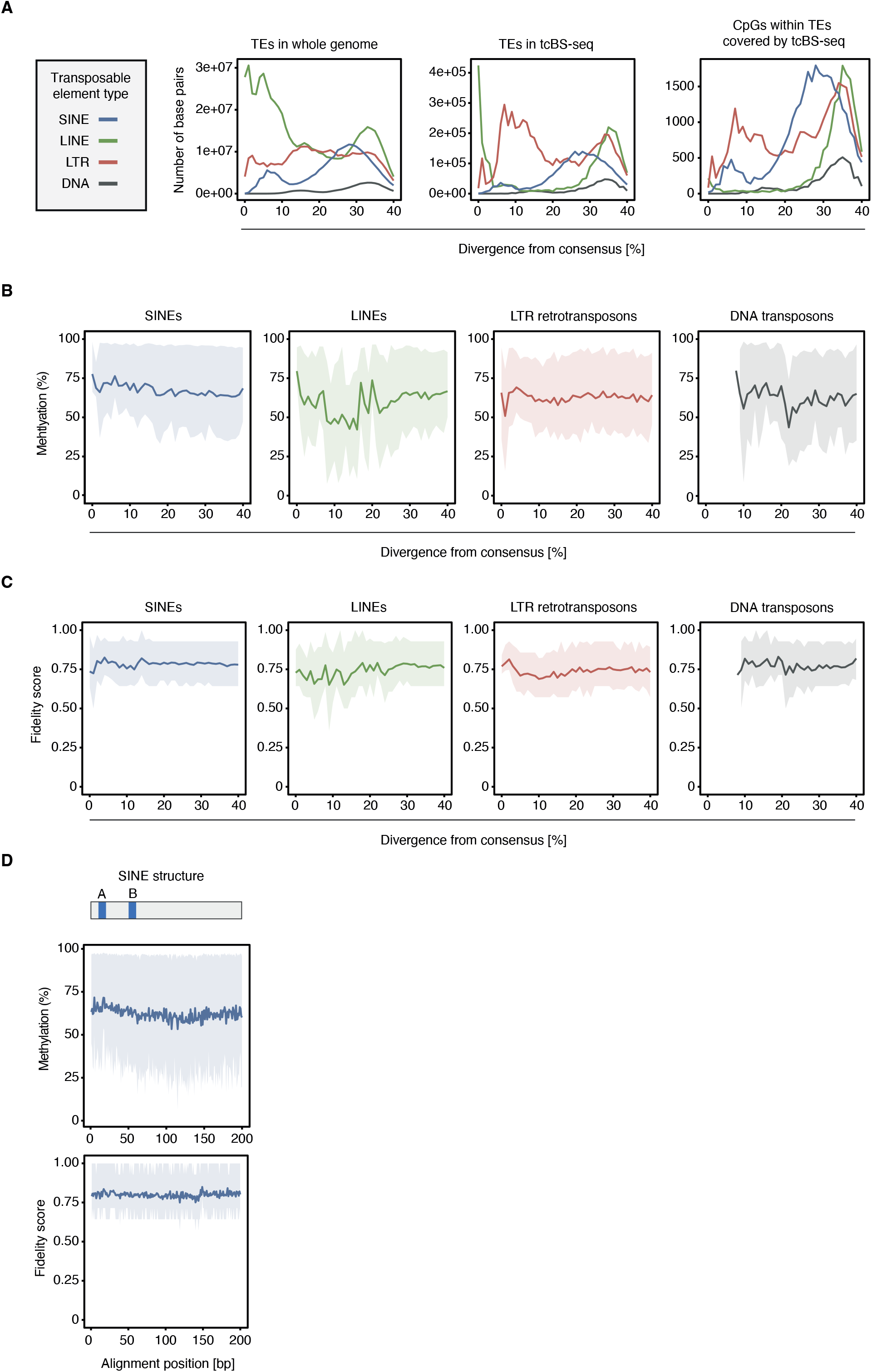
Methylation fidelity at transposable elements. (A) Plots showing base pair counts for TEs in the whole genome (left), TEs represented by the tcBS-seq data (middle), and the CpGs within TEs represented by the tcBS-seq data, with varying percent divergences from the consensus sequence. TE types are represented by colour: SINEs in blue; LINEs in green; LTR retrotransposons (LTR) in red; DNA transposons (DNA) in grey. (B and C) Plots of methylation level (B) or fidelity score (C) versus percent divergence from the consensus sequence for different TE types. (D) Plots of methylation level and fidelity score versus the position of a CpG within a SINE. Top bar shows the archetypal SINE structure relative to the alignment position on the x-axis of the plots below, with A- and B-block promoter regions coloured in blue. For all plots, methylation levels and fidelity score are calculated as averages across CpGs in each TE; lines represent mean values, while ribbons show the interquartile range.

**Fig. S8:**
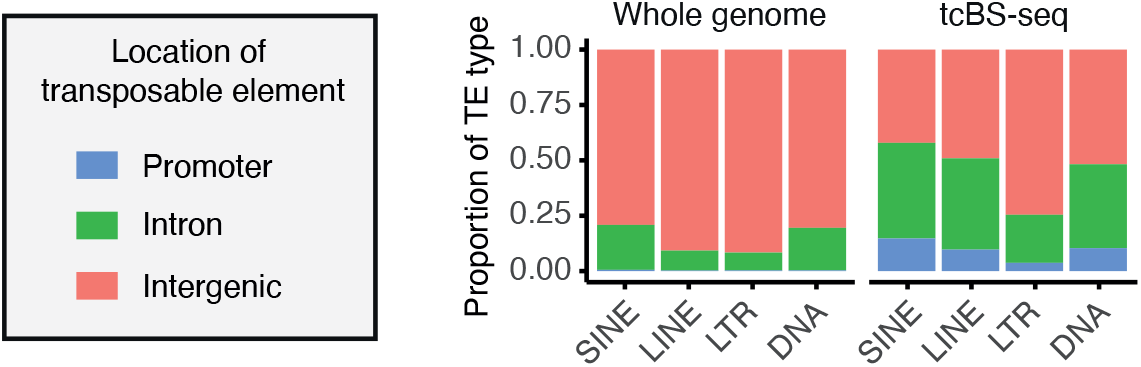
Enrichment of genically located transposable elements in tcBS-seq. Proportion of TE types across three different genomic locations (intergenic, intron, and promoter) both genome-wide (left) and in the tcBS-seq data (right). Genomic locations are represented by colour: promoter in blue; intron in green; intergenic in orange.

**Fig. S9:**
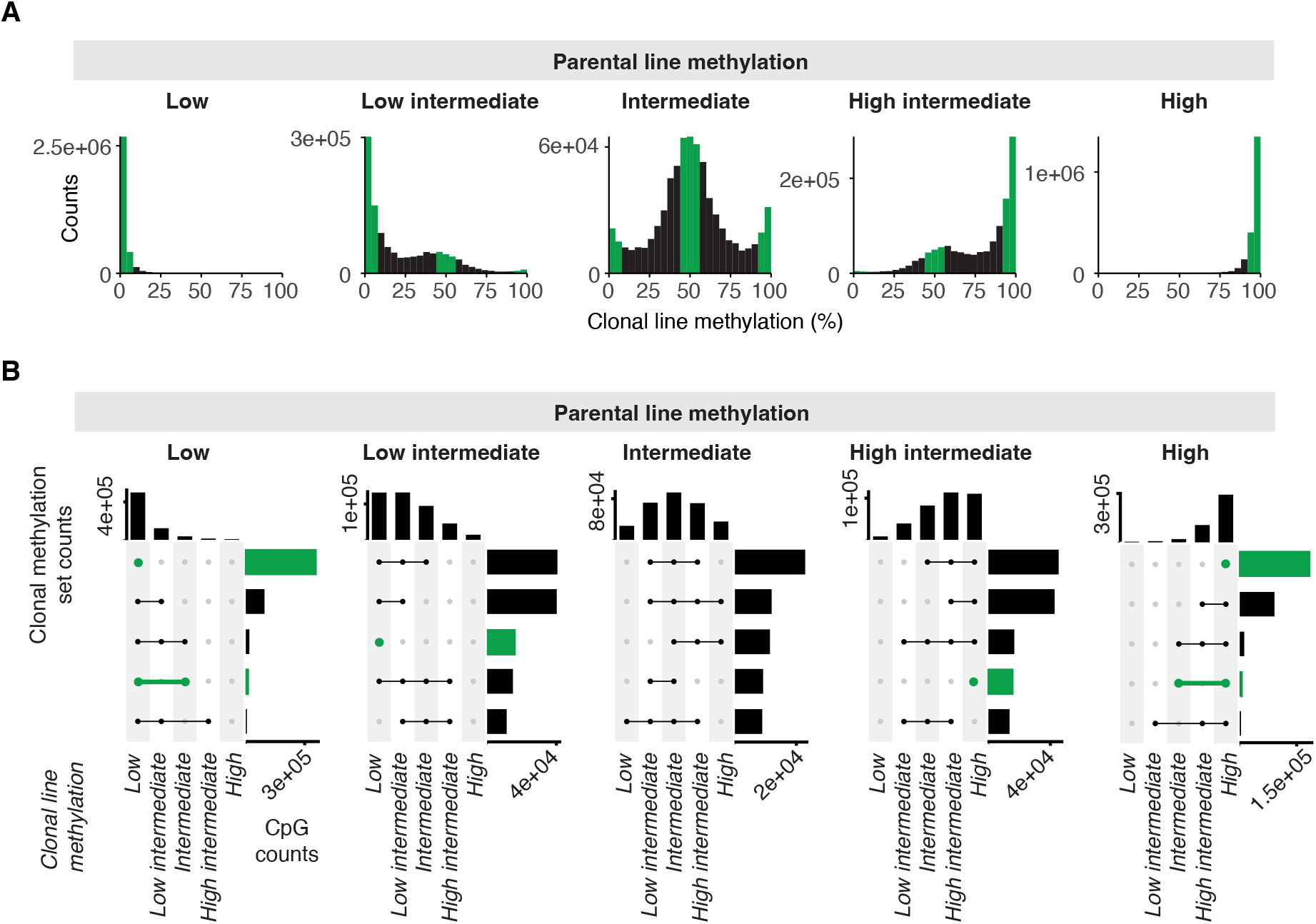
Intermediately methylated CpGs are prone to probabilistic inheritance between cell divisions in MEF-2 cell lines. (A) Clonal line methylation distributions from different parental line methylation states for the MEF-2 cell lines. (B) UpSet plots of clonal line methylation states per CpG from different parental line methylation states for the MEF-2 cell lines. In each panel, the top bar graph shows the counts of distinct clonal methylation states per CpG. The circles in each panel represent the clonal methylation states per CpG, where multiple states per CpG are represented by circles connected by a line - the attached horizontal bar graph shows the CpG counts that exhibit a corresponding combination of clonal methylation states shown by the circles. Only the five most representative clonal methylation state combinations are shown. Green bars represent cases of potential faihtful methylation inheritance because this kind of methylation inheritance will only result in 0, 50, or 100% methylation states in the clonal lines. Low = 0-10%, Low intermediate = 10-40%, Intermediate = 40-60%, High intermediate = 60-90%, High = 90-100% methylation.

**Fig. S10:**
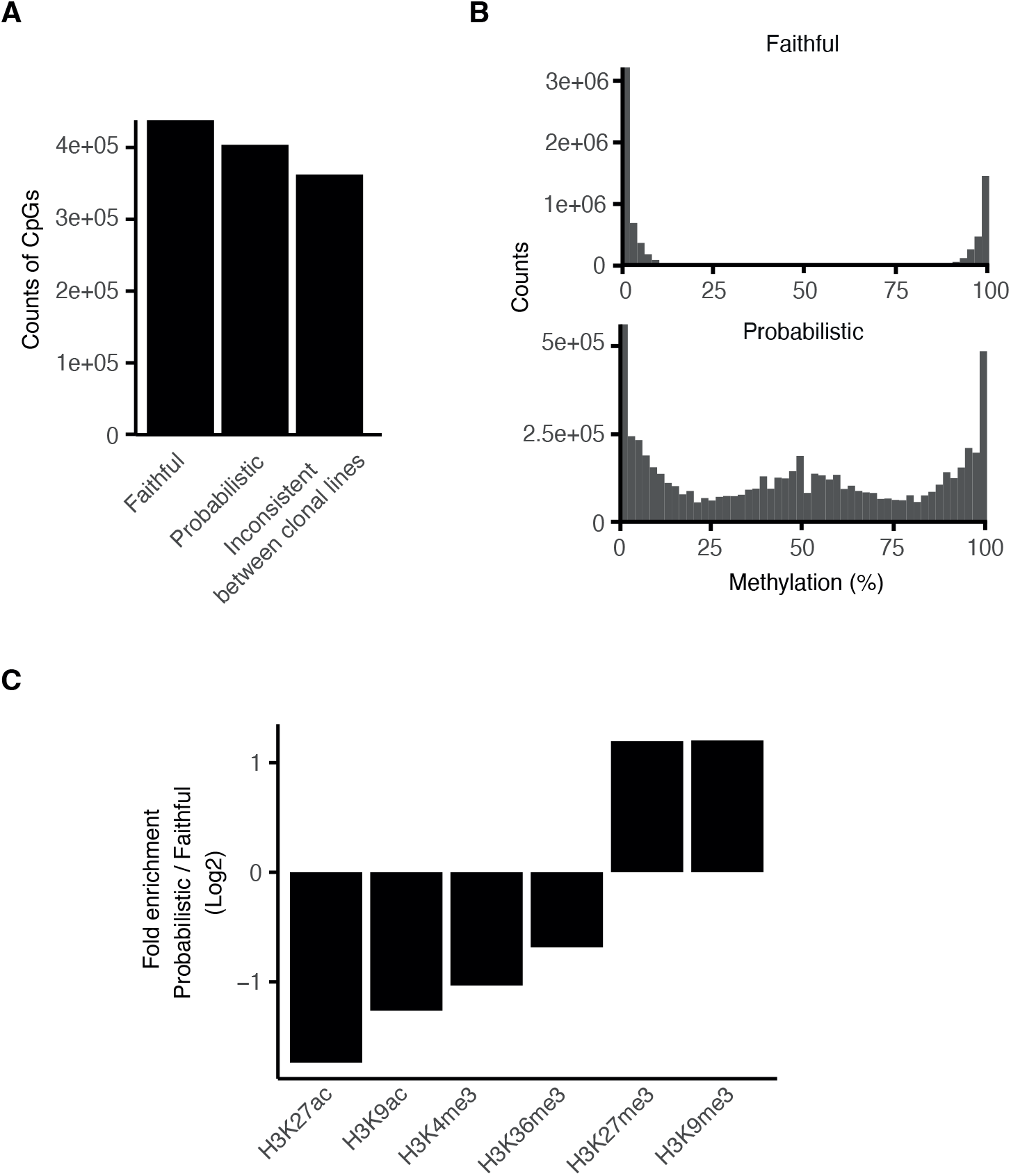
Probabilistically methylated CpGs associate with repressive histone tail modifications H3K27me3 and H3K9me3. A) Counts of faithfully and probabilistically methylated CpGs. CpGs are defined as faithful if all clonal lines exhibit [0-10], (40-60], or (90-100] % methylation. If at least one clonal line exhibits (10-40] or (60-90] % methylation in both MEF-1 and MEF-2, we define the CpGs as probabilistically methylated. CpGs that are inconsistently characterised as faithful or probabilistic between MEF-1 and MEF-2 are not considered for further analyses. (B) Methylation distributions of faithful and probabilistic CpGs across all MEF-1 and MEF-2 cell lines. (C) Enrichment of histone tail modification peak overlap with probabilistic versus faithful CpGs.

**Fig. S11:**
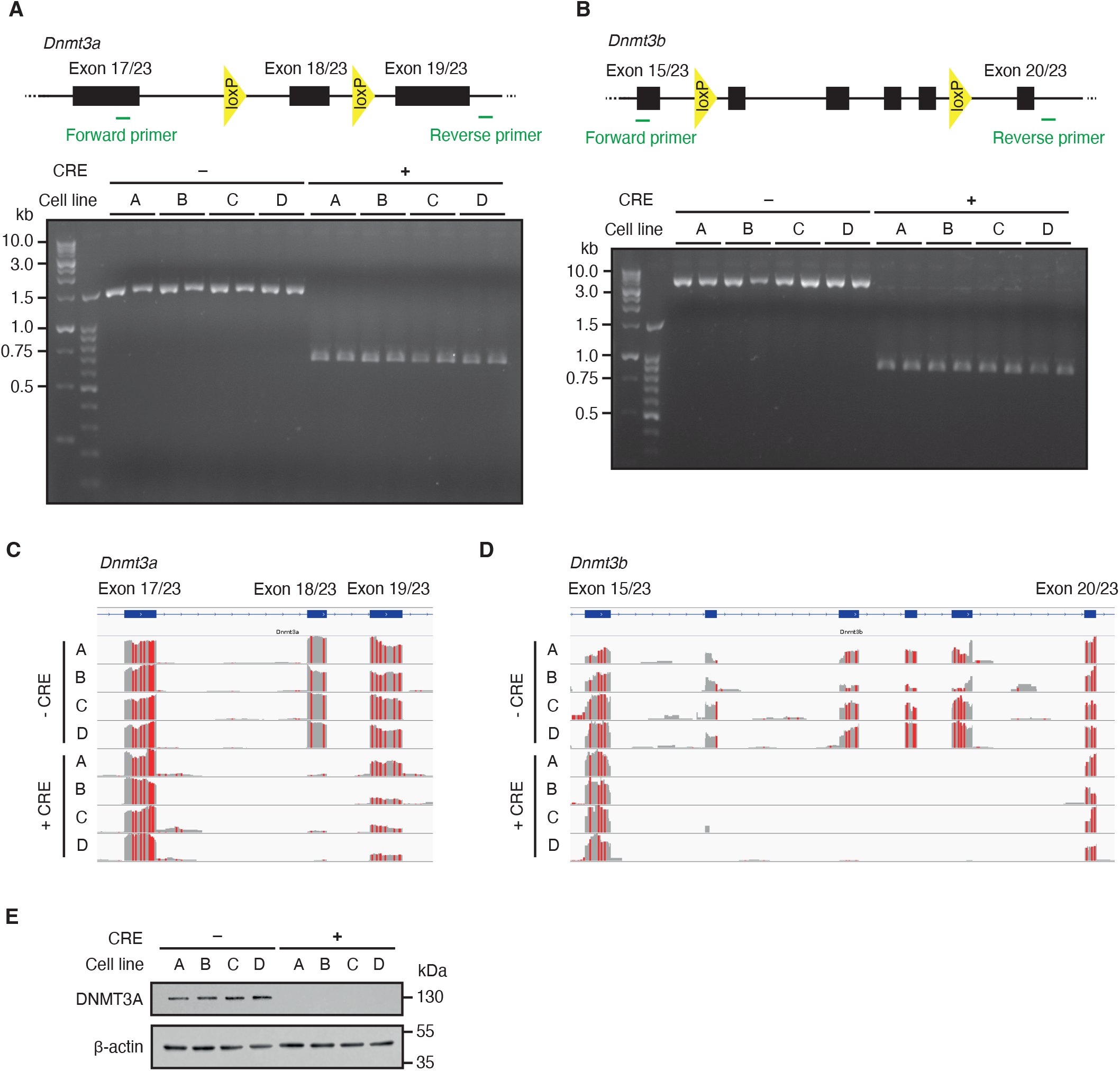
Confirmation of induced Dnmt3a/3b DKO in primary MEFs. (A) Top: diagram showing the location of the loxP sites flanking exon 18 of *Dnmt3a*, as well as the location of the primers used to confirm the knockout (KO). Bottom: agarose gel image of a PCR to confirm the TAT-CRE induced KO of *Dnmt3a* in the DNA. (B) Top: diagram showing the location of the loxP sites flanking exons 16-19 of *Dnmt3b*, as well as the location of the primers used to confirm the KO. Bottom: agarose gel image of a PCR to confirm the TAT-CRE induced KO of *Dnmt3b* in the DNA. (C and D) IGV screenshots of RNA-seq data showing the absence of transcripts that include the loxP flanked exons for both *Dnmt3a* and *Dnmt3b*. (E) Western blots of untreated and TAT-CRE-treated cell lines for DNMT3A and loading control ß-actin.

**Table S1:**
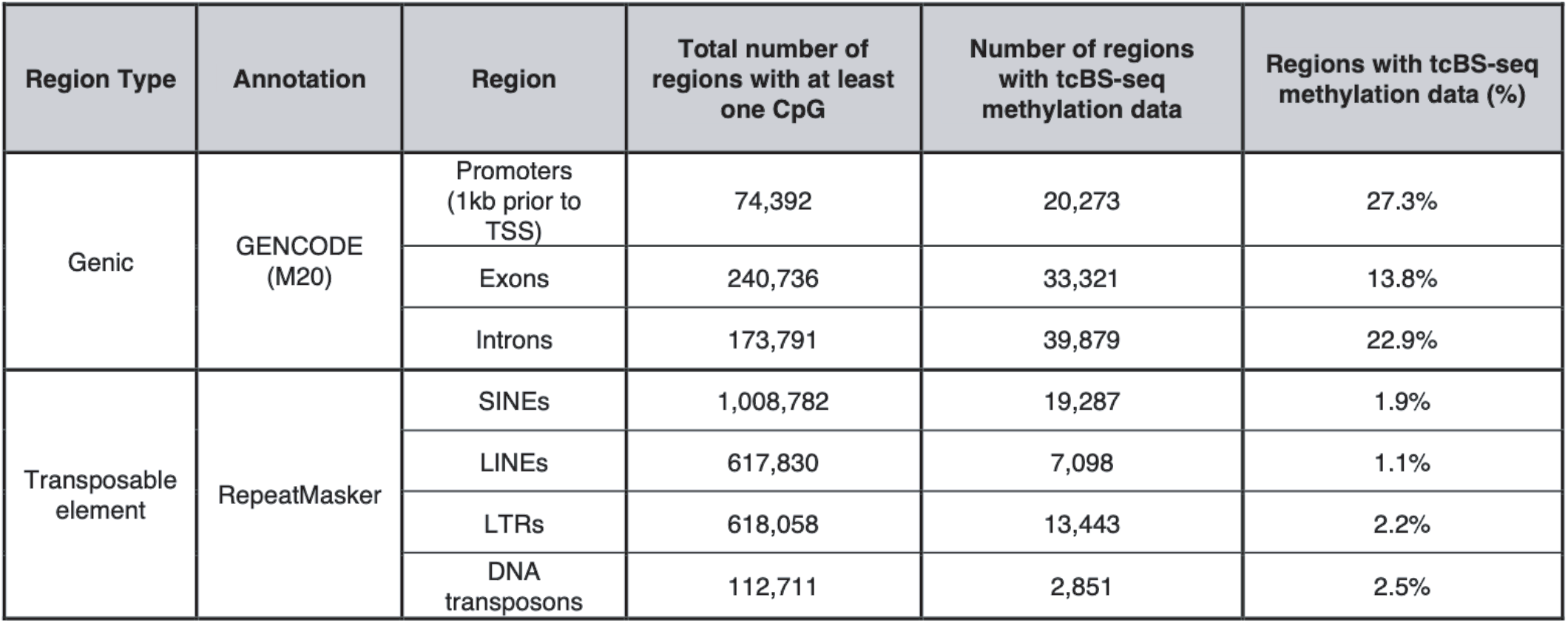
Coverage of genic regions and transposable elements by target capture bisulphite sequencing.

**Table S2:** Publicly available bisulfite-sequencing and RNA-sequencing datasets.

| Cell type | Library Prep Type | GEO / ENCODE accession | ENCODE (Bed file) |
| --- | --- | --- | --- |
| Primary MEFs | WGBS | GSM2648457 | - |
| 3T3 immortalised MEFs | WGBS | GSM4942823, GSM4942824, GSM4942825 | - |
| E13.5 mouse limb | WGBS | ENCSR700FCF | ENCFF556ENA, ENCFF834PXN |
| E14.5 mouse limb | WGBS | ENCSR099RYD | ENCFF176VLZ, ENCFF038IVS |
| mESCs<br>(maintained with 2i + LIF, and established with low PD) | WGBS | GSM2425451, GSM2425460 | - |
| mESCs<br>(maintained with 2i + LIF) | WGBS | GSM1027570 | - |
| mESCs<br>(maintained with serum) | WGBS | GSM1027571 | - |
| mESCs<br>(grown on feeder cells) | tcBS-seq | GSM3713369, GSM3713370, GSM3713371 | - |
| E3.5 inner cell mass | WGBS | GSM2229982, GSM2229983 | - |
| mESCs<br>(maintained with 2i + LIF, and established with low PD) | poly-A RNA-seq | GSM2425493, GSM2425494 | - |

**Table S3:**
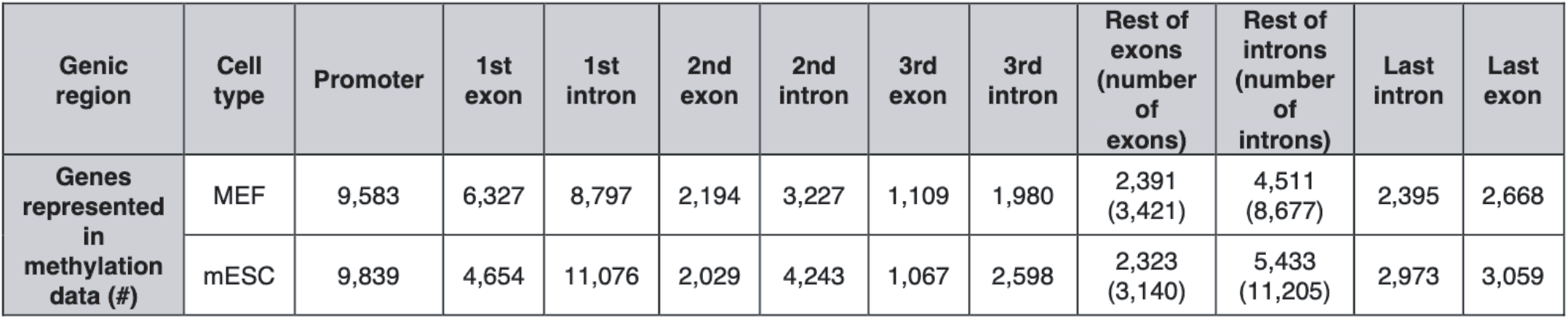
Number of genes and genic regions represented in methylation data.

**Table S4:**
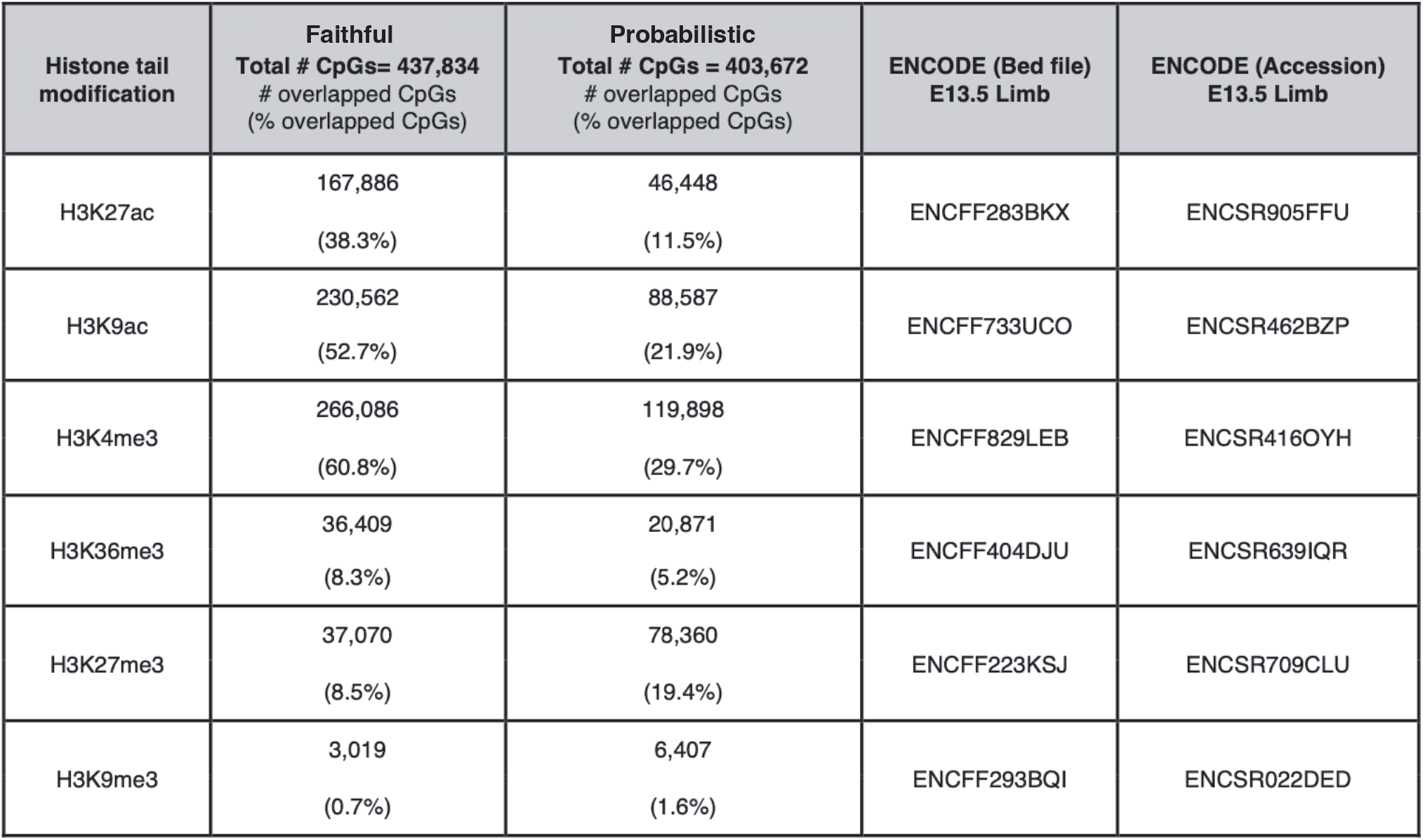
Counts and ratios of faithful and probabilistic CpG overlap with various histone tail modification peaks.

| Histone tail modification | Faithful<br>Total # CpGs= 437,834<br># overlapped CpGs<br>(% overlapped CpGs) | Probabilistic<br>Total # CpGs = 403,672<br># overlapped CpGs<br>(% overlapped CpGs) | ENCODE (Bed file)<br>E13.5 Limb | ENCODE (Accession)<br>E13.5 Limb |
| --- | --- | --- | --- | --- |
| H3K27ac | 167,886<br>(38.3%) | 46,448<br>(11.5%) | ENCFF283BKX | ENCSR905FFU |
| H3K9ac | 230,562<br>(52.7%) | 88,587<br>(21.9%) | ENCFF733UCO | ENCSR462BZP |
| H3K4me3 | 266,086<br>(60.8%) | 119,898<br>(29.7%) | ENCFF829LEB | ENCSR416OYH |
| H3K36me3 | 36,409<br>(8.3%) | 20,871<br>(5.2%) | ENCFF404DJU | ENCSR639IQR |
| H3K27me3 | 37,070<br>(8.5%) | 78,360<br>(19.4%) | ENCFF223KSJ | ENCSR709CLU |
| H3K9me3 | 3,019<br>(0.7%) | 6,407<br>(1.6%) | ENCFF293BQI | ENCSR022DED |

